# Analysis of spike protein variants evolved in a novel mouse model of persistent SARS-CoV-2 infection

**DOI:** 10.1101/2023.03.19.533317

**Authors:** Dongbum Kim, Jinsoo Kim, Minyoung Kim, Heedo Park, Sony Maharjan, Kyeongbin Baek, Bo Min Kang, Suyeon Kim, Sangkyu Park, Man-Seong Park, Younghee Lee, Hyung-Joo Kwon

**Author notes:** Correspondences: Man-Seong Park, Department of Microbiology, College of Medicine, and the Institute for Viral Diseases, Korea University, Seoul, 02841, Republic of Korea.; Younghee Lee, Department of Biochemistry, College of Natural Sciences, Chungbuk National University, Cheongju 28644, Republic of Korea.; Hyung-Joo Kwon, Department of Microbiology, College of Medicine, Hallym University, Chuncheon 24252, Republic of Korea. These authors contributed equally: Dongbum Kim, Jinsoo Kim, Minyoung Kim, Heedo Park.

## Abstract

SARS-CoV-2 mutation rates have increased over time, resulting in the emergence of several variants of concern. Persistent infection is assumed to be involved in the evolution of the variants; however, there is currently no animal model to recapitulate persistent infection. We established a novel model of persistent infection using xenografts of Calu-3 human lung cancer cells in immunocompromised mice. After infection with wild-type SARS-CoV-2, viruses were found in the tumor tissues for up to 30 days and acquired various mutations, predominantly in the spike (S) protein, some of which increased while others fluctuated for 30 days. Three isolated viral clones with defined mutations produced higher virus titers than the wild-type virus in Calu-3 cells without cytotoxic effects. In K18-hACE2 mice, the variants were less lethal than the wild-type virus. Infection with each variant induced production of cross-reactive antibodies to the receptor binding domain of wild-type S protein and provided protective immunity against subsequent challenge with wild-type virus. These results suggest that most of the SARS-CoV-2 variants acquired mutations promoting host adaptation in the Calu-3 xenograft mice. This model can be used in the future to further study persistent SARS-CoV-2 infection.

## Introduction

Severe acute respiratory syndrome coronavirus-2 (SARS-CoV-2) is the highly contagious novel coronavirus responsible for the coronavirus disease 2019 (COVID-19) pandemic^1^. As of March 07, 2023, the World Health Organization reported over 759 million infections and more than six million deaths due to COVID-19 (https://covid19.who.int). Despite various countermeasures and global vaccination efforts to lower SARS-CoV-2 transmission, there is still a significant risk of further COVID-19 waves resulting from the emergence of novel SARS-CoV-2 variants^2^.

The RNA genome of SARS-CoV-2 is similar to that of other SARS-related coronaviruses, encoding four major structural proteins, referred to as the spike (S), membrane (M), nucleocapsid (N), and envelope (E), and a variable number of open reading frames (ORFs) that assist in viral replication and transcription^3^. The receptor binding domain (RBD) of the S protein facilitates viral entry into host cells via its high affinity to human angiotensin-converting enzyme 2 (ACE2)^4^. Two other highly pathogenic coronaviruses, SARS-CoV and Middle East respiratory syndrome coronavirus (MERS-CoV), caused global pandemics with high mortality in 2002 and 2012, respectively^1^. Although the pandemics caused by SARS-CoV and MERS-CoV each lasted less than a year, the COVID-19 pandemic has lasted over 3 years, raising fundamental issues regarding the frequent mutation and evolutionary patterns of viral genes^5^. Specifically, increased mutation seems to influence virus transmissibility, pathogenicity, infectivity, immune escape, drug resistance, and antigenicity^6, 7^.

Shortly after the COVID-19 outset, a SARS-CoV-2 variant harboring a D614G mutation in the S protein rapidly spread as a dominant strain, and more variants have since emerged^8, 9^. Successively emerging strains of SARS-CoV-2 have been labeled as Alpha, Beta, Gamma, Delta, Epsilon, Eta, Iota, Kappa, Mu, Zeta, and Omicron variants. The Omicron variant has been classified as a variant of interest, whereas the Alpha, Beta, Gamma, and Delta variants have been classified as variants being monitored (https://www.cdc.gov/coronavirus/2019-ncov/variants/variant-classifications.html). Omicron is currently the predominant variant worldwide, accounting for > 99% of the variant genome frequency as of March 2023 (https://gisaid.org/hcov19-variants/). Compared with the previous variants, the Omicron variant contains 30 or more mutations in the S protein^10^.

Individuals that are immunocompromised because of cancer, chemotherapy, organ transplantation, HIV infection, chronic diabetes, or other reasons cannot neutralize SARS-CoV-2, so they sustain persistent infection for long periods (e.g., over 3 months in a patient with chronic lymphoid leukemia)^11–14^. When viruses multiply inside human tissues for long periods, they can evolve adaptations to evade host immune responses and resist the effects of antiviral drugs, increasing the probability that a novel strain will emerge. Hence, immunocompromised individuals represent a likely source of new SARS-CoV-2 variants^12, 13^. There is currently no animal model to recapitulate persistent SARS-CoV-2 infection. Previously, we reported that cellular responses after SARS-CoV-2 infection, especially apoptosis, differ depending on the cell type^15^. Vero monkey kidney cells show high virus production and an apoptotic phenotype after infection, whereas Calu-3 human lung cancer cells persistently grow without apoptosis and release low virus titers after infection. We postulated that the response of Calu-3 cells mimics persistent SARS-CoV-2 infection in humans. In the present study, we created xenograft tumors derived from Calu-3 cells in immunocompromised mice and infected them with SARS-CoV-2. We found that new SARS-CoV-2 variants emerged in the tumor tissues, which we characterized *in vitro* and *in vivo.* Our results suggest that the mouse xenograft model can be useful for further investigations of persistent SARS-CoV-2 infection.

## Results

### Establishment of a persistent infection model using lung cancer xenografts in mice

To test our hypothesis that Calu-3 cells can be used in mice to recapitulate persistent SARS-CoV-2 infection, we first induced xenograft tumors into genetically immunocompromised NOD/ShiLtJ-Rag2*^em1AMC^*Il2rg*^em1AMC^*(NRGA) mice using Calu-3 cells. We then infected the xenograft tumors with wild-type SARS-CoV-2 and monitored the mice at 3, 15, and 30 dpi as depicted in Fig. 1a. The infected mice showed similar body weight, tumor volume, and tumor weight compared to uninfected controls, suggesting that virus infection did not induce prominent side effects (Fig. 1b). Importantly, for up to 30 dpi, more than 10^4^ pfu of virus were found in the tumors but not the sera, lungs, or brains of the mice (Fig. 1c). As shown in Fig. 1d, expression of the viral N protein increased during the course of infection, suggesting persistent and increasing virus replication in the tumors. Immunohistochemistry data revealed persistent expression of the viral S and N proteins along with hACE2 in the tumor tissues (Fig. 1e). When the same strategy was applied to A549 cells, another lung cancer cell line, virus was found at 3 and 15 dpi but not 30 dpi, probably because the A549 cells had little hACE2 expression and different cellular responses compared with the Calu-3 cells^16^ (Supplementary Data Fig. 1). Taken together, our results confirmed that the xenograft model established with highly permissible Calu-3 lung cancer cells can be used as a model for persistent SARS-CoV-2 infection without harmful effects on infected mice.

**Fig. 1.**
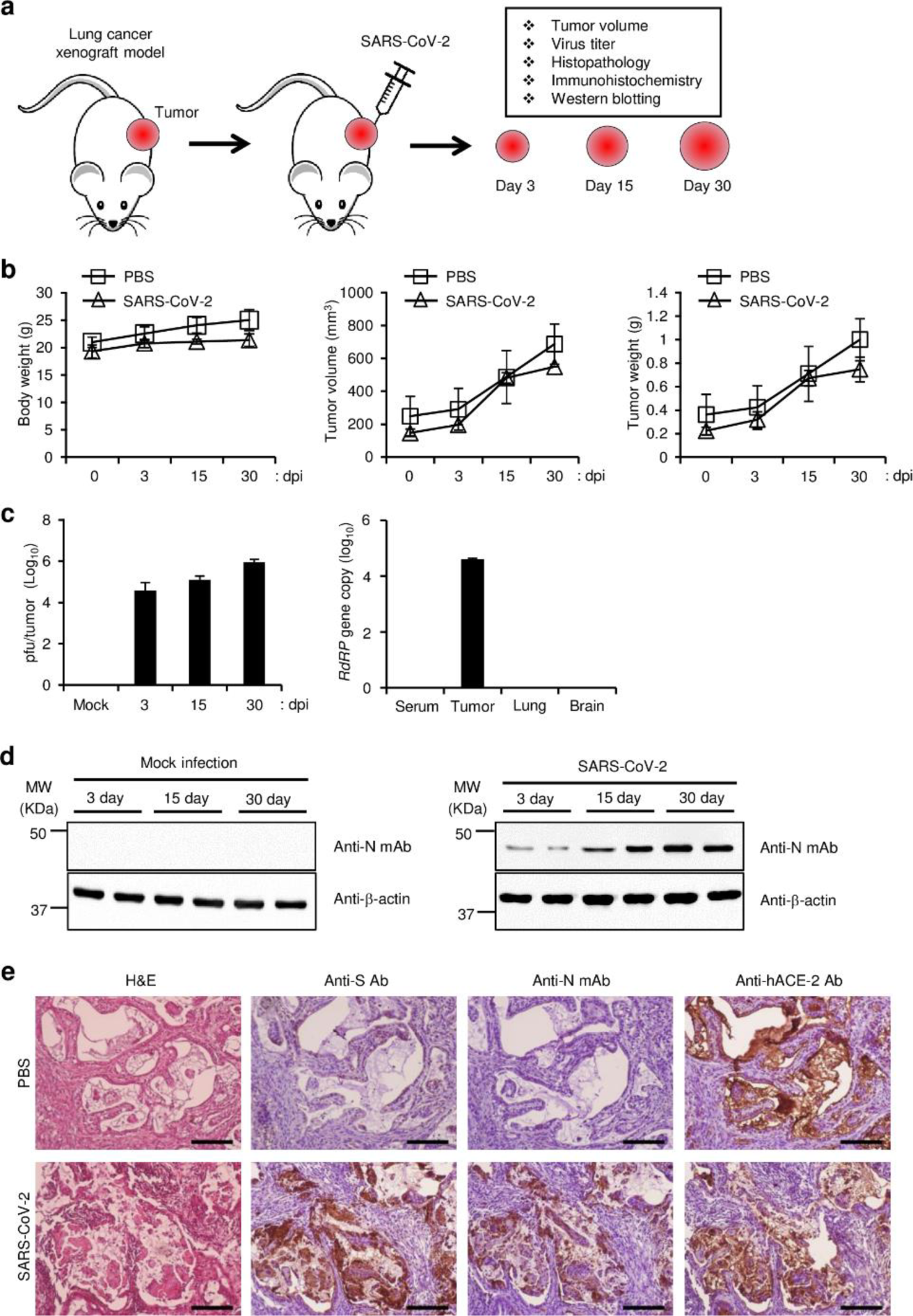
Replication of SARS-CoV-2 in a lung tumor xenograft mouse model. **a,** Schematic diagram of the experimental procedure. Calu-3 cells were subcutaneously implanted into the right flank of NRGA mice (n = 3 per group). After the tumor volume reached 100 mm^3^, the tumors were inoculated with 1 × 10^6^ pfu of wild-type SARS-CoV-2, and the virus titers in the tumors were analyzed at 3, 15, and 30 dpi. **b,** Body weight of each infection group (left panel). Tumor volume of each infection group (middle panel). Tumor weight of each infection group (right panel). Data are presented as mean ± s.d. **c,** Replication of SARS-CoV-2 in lung tumors. Tumor tissues were collected at 3, 15, and 30 dpi. Viral titers in the supernatants of tumor homogenates were determined by plaque assay (left panel). Serum, tumor, lung, and brain were collected at 30 dpi, and viral titers were determined by qRT-PCR analysis for the SARS-CoV-2 *RdRP* gene (right panel). Data are presented as mean ± s.d. **d,** Expression of SARS-CoV-2 N protein in tumor tissues. The expression levels of SARS-CoV-2 N protein in mock-infected and SARS-CoV-2-infected tumor homogenates were determined by western blot. Expression of β-actin was used as a control. **e,** Expression of SARS-CoV-2 N protein and S protein in tumor tissues. Mice (n=2) were sacrificed at 30 dpi. Paraformaldehyde-fixed, paraffin-embedded tumor tissues were sliced to 5 µm thickness, and H&E staining and immunohistochemical staining were performed to detect SARS-CoV-2 S protein and N protein and hACE2. Scale bars, 25 µm. dpi, days post-infection; Anti-N mAb, anti-SARS-CoV-2 N protein monoclonal antibody; Anti-S Ab, anti-SARS-CoV-2 S protein polyclonal antibody.

### Appearance of SARS-CoV-2 variants in the mouse xenograft model

To investigate whether persistent SARS-CoV-2 infection in the mouse xenograft model results in the occurrence of variants, we first expanded the original wild-type virus stock in three passages of cell culture and used the sample as a start control (passage 3). We then infected the tumors of three mice with wild-type SARS-CoV-2 and collected tumor tissues at 3, 15, and 30 dpi. We synthesized viral cDNA from the supernatants of tumor homogenates and analyzed the sequences by whole-genome sequencing. Compared with the wild-type virus, the passage 3 viruses and the viruses from the xenograft tumors had a total of six mutations at five residues in the S protein and one mutation each in ORF1ab, ORF3a, the M protein, ORF6, and ORF8 (Fig. 2). We focused on the mutations in the S protein, which are shown in Supplementary Data Table 1. The mutations T95I, D178N, E484D, H655Y, and R685S were found in both the passage 3 viruses and the viruses from the tumor tissues. The mutations N148K, E180G, N185K, and D215G were found only in the viruses from the tumor tissues, whereas the mutation D178G was found only in the passage 3 viruses.

**Fig. 2.**
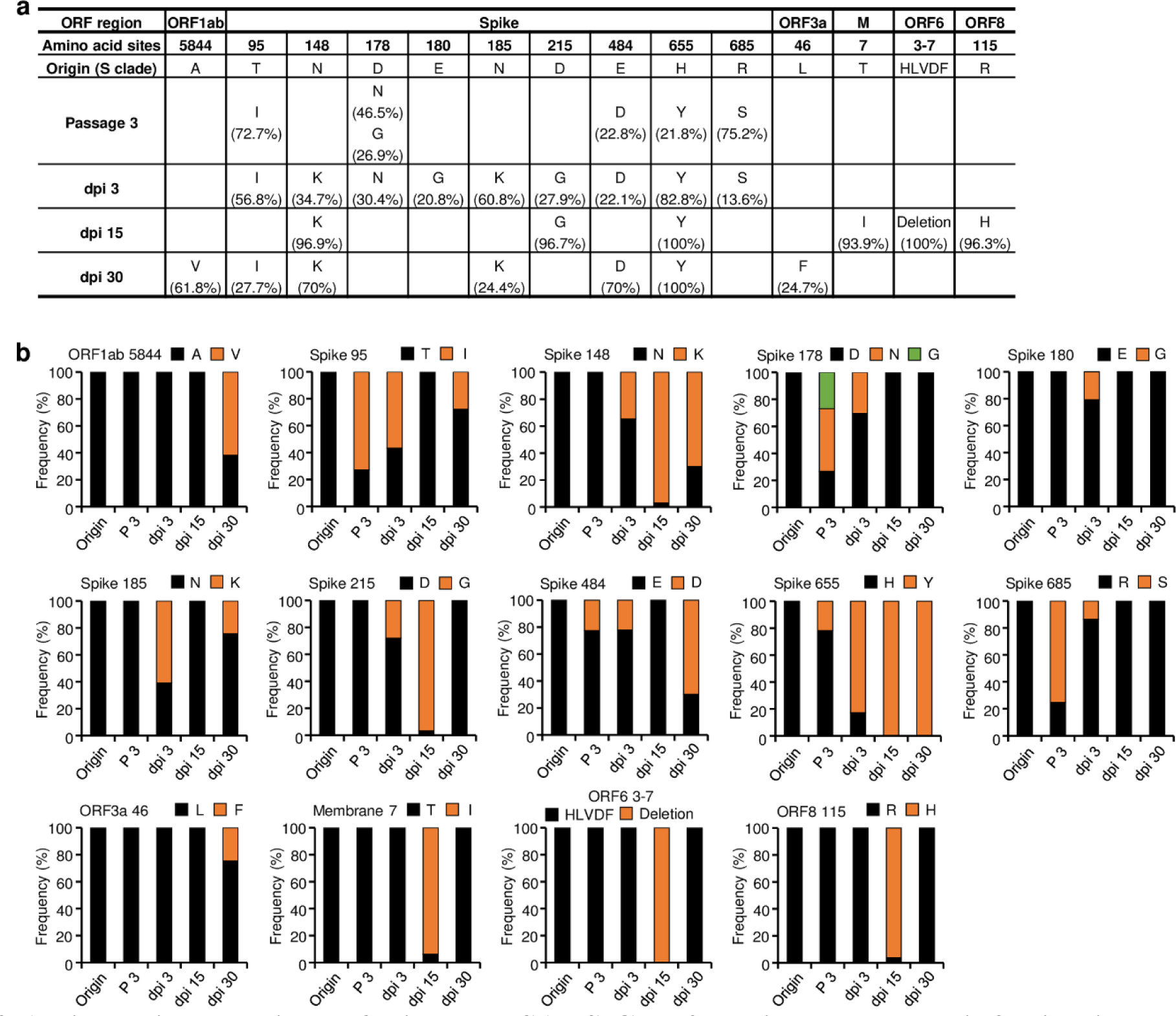
Amino acid mutations of wild-type SARS-CoV-2 during long-term infection in lung tumor tissue. Calu-3 cells were subcutaneously implanted into the right flank of NRGA mice (n=1 per group). After the tumor volume reached 100 mm^3^, the tumors were inoculated with 1 × 10^6^ pfu of wild-type SARS-CoV-2. Whole-genome sequences of viruses in the tumors at 3, 15, and 30 dpi were analyzed. **a,** Frequency of mutations in wild-type SARS-CoV-2 during long-term infection in lung tumor tissue. **b,** Frequencies and substitutions of amino acid sequences in wild-type SARS-CoV-2 during long-term infection in lung tumor tissue. Origin (wild type): amino acid sequences of SARS-CoV-2 wild type (Table S1). Passage 3 (P 3): amino acid sequences in wild-type SARS-CoV-2 were analyzed after amplification in three passages in Vero E6 cells. dpi, days post-infection.

Most of the mutations in the S protein were previously observed in clinically important SARS-CoV-2 variants or in SARS-CoV-2 derived from cell culture^17–22^(https://coVariants: Shared Mutations, https://gisaid.org/hcov19-mutation-dashboard GISAID). The mutations T95I, D178G, D215G, and H655Y were reported previously in various variants^17–20^(https://coVariants: Shared Mutations). The mutation E484D occurred at the same residue as the mutations E484K, E484Q, and E484A^17^(https://coVariants: Shared Mutations) that were previously found in cultures of human airway H522 cells^20^. The mutation R685S was previously isolated from VSV-GFP-SARS-CoV-2-S_Δ21_ during propagation in MA104 cells^20^. Although the frequencies of most of the mutations fluctuated over time in the mice, the frequency of mutation H655Y approached 100%, and the mutations N148K and E484D were maintained at high frequencies at 30 dpi. Some mutations transiently appeared in the mice but were later displaced by the original wild-type sequences. There were also a few fluctuating mutations in ORF1ab, ORF3a, M, ORF6, and ORF8. Considering that various mutations, including reportedly significant ones in the S protein, arose and were maintained in the mouse xenograft model for up to 30 days, it is likely that the model successfully recapitulates the evolution of variants during persistent infection^11–14^.

### Isolation and characterization of viral clones from tumors infected with wild-type SARS-CoV-2

Next, we analyzed the S protein sequences of individual viruses obtained from the mouse xenografts at 30 dpi. As most of the confirmed mutations were located on the S1 subunit of the S protein, we cloned the cDNA sequences of that region and analyzed them by further DNA sequencing (Supplementary Data Fig. 2a). As shown in Supplementary Data Fig. 2b, the sequencing of the S1 subunit revealed three different variants (variants 1, 2, and 3) based on the combinations of mutations across six amino acid residues. The D178N mutation was not found in the whole-genome sequence analysis of tumor-derived viruses at 30 dpi (Fig. 2), suggesting that this mutation was a minor variation that fell under the cutoff level for detection in the whole-genome analysis.

To specifically characterize SARS-CoV-2 variants that emerged in the xenograft tumors, we isolated 15 viral clones from individual plaques after plaque formation assays with tumor homogenates obtained at 30 dpi. Then, we identified three clones (S-1, S-6, and S-9) corresponding to the three variants based on the S1 subunit sequences and performed whole-genome sequencing on the clones. Because we expanded the three clones with three passages in Vero E6 cells, we also analyzed the wild-type virus after cell culture expansion as a control (passages 3 and 6). The results showed that the three clones had the expected S mutations along with a mutation in the N protein and three mutations in the ORF1ab (Fig. 3). Considering that the wild-type viral sequences changed during culture (passage 3 vs. passage 6), we cannot exclude the possibility that the exact sequences of the clones initially obtained from the mouse xenografts differed from the sequences after three passages. The expanded clones from the xenografts and the wild-type pool from cell culture both included a mixture of genotypes at our residues of interest; however, all three expanded clones had the H655Y mutation at or near 100% frequency, no presence of the S685 mutation, and the N protein mutation T24I at a frequency of at least 30%.

**Fig. 3.**
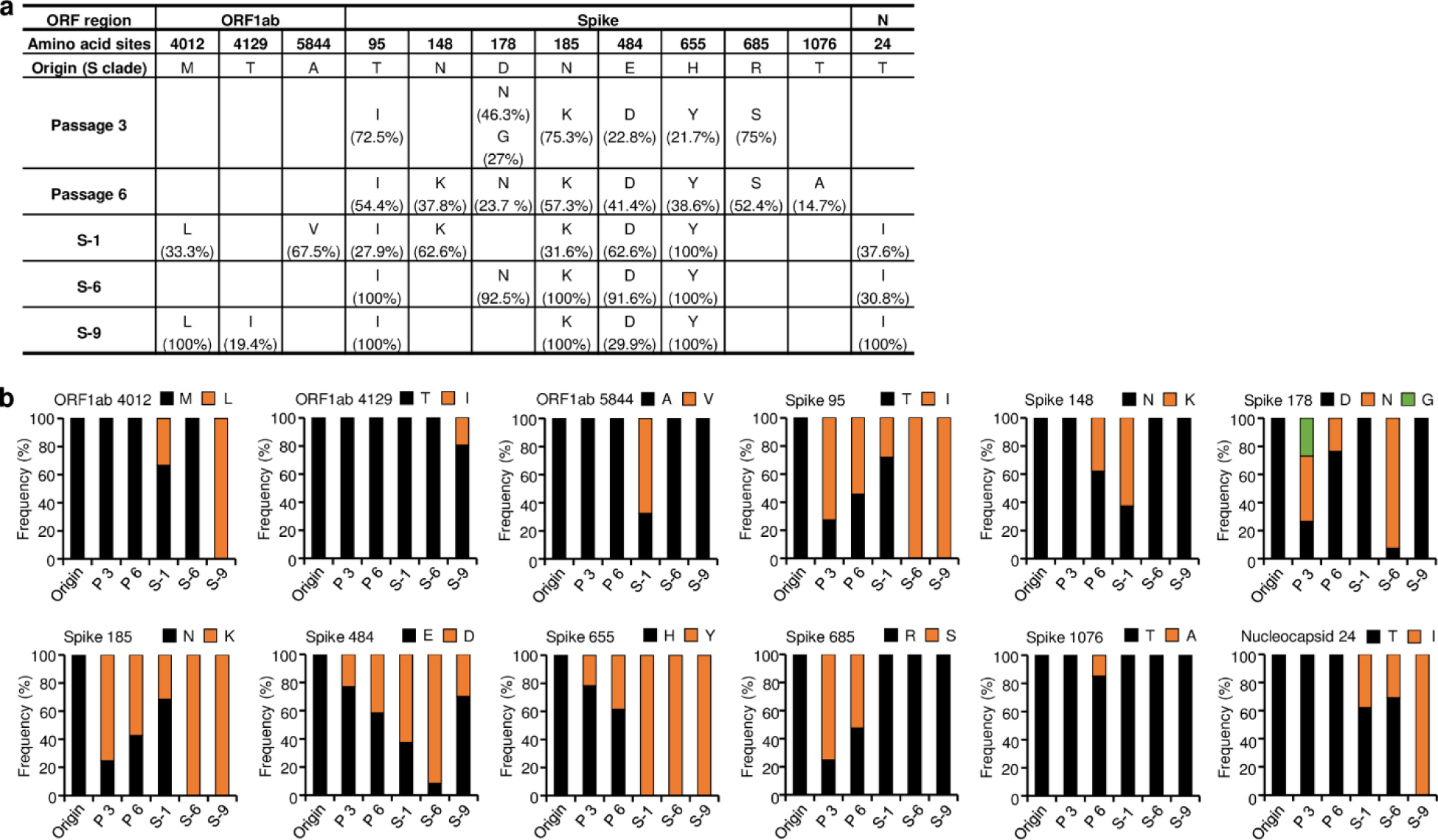
Viral clone isolation from plaques and analysis of amino acid mutations. Viral clones were isolated at 30 dpi from plaques derived from supernatants of tumors infected with wild-type SARS-CoV-2. Fifteen viral clones were amplified in three passages in Vero E6 cells. cDNA was synthesized from each clone, and the S1 subunit region of the S protein was further cloned and analyzed by DNA sequencing. Three selected viral clones (S-1, S-6, S-9) corresponding to the three types of variants in Figure S2 were collected, and their whole-genome sequences were analyzed. **a,** Comparison of mutations in the three viral clones and the origin (wild type). **b,** Accumulations and substitutions of amino acid sequences in the three viral clones. Origin (wild type): amino acid sequences of wild-type SARS-CoV-2 (Supplementary Table 1). Passage 3 (P 3) and Passage 6 (P 6): amino acid sequences in wild-type SARS-CoV-2 after amplification in three and six passages, respectively, in Vero E6 cells. dpi, days post-infection.

To further compare the clones isolated from the xenografts with the original wild-type, we investigated cytotoxic effects of the clones in Vero E6 cells and Calu-3 cells. In Vero cells, the wild-type virus and clones S-1, S-6, and S-9 induced clear cytotoxic effects to a similar extent (Fig. 4a,b). However, no cytotoxic effects were induced by any of the viruses in Calu-3 cells (Fig. 4c), suggesting that Calu-3 cells, but not Vero cells, recapitulate persistent SARS-CoV-2 infection without apoptosis as we previously reported^15^. When we measured virus replication 48 h after infection by plaque assay, there was no difference between the wild type and the three clones in Vero E6 cells; however, all three clones showed higher virus titers than the wild type in Calu-3 cells (Fig. 4d). Considering that the wild-type virus replicates less in Calu-3 cells than in Vero E6 cells, the effects of mutation could be monitored more easily in Calu-3 cells. It is likely that the mutations contributed to the increased replication in Calu-3 cells.

**Fig. 4.**
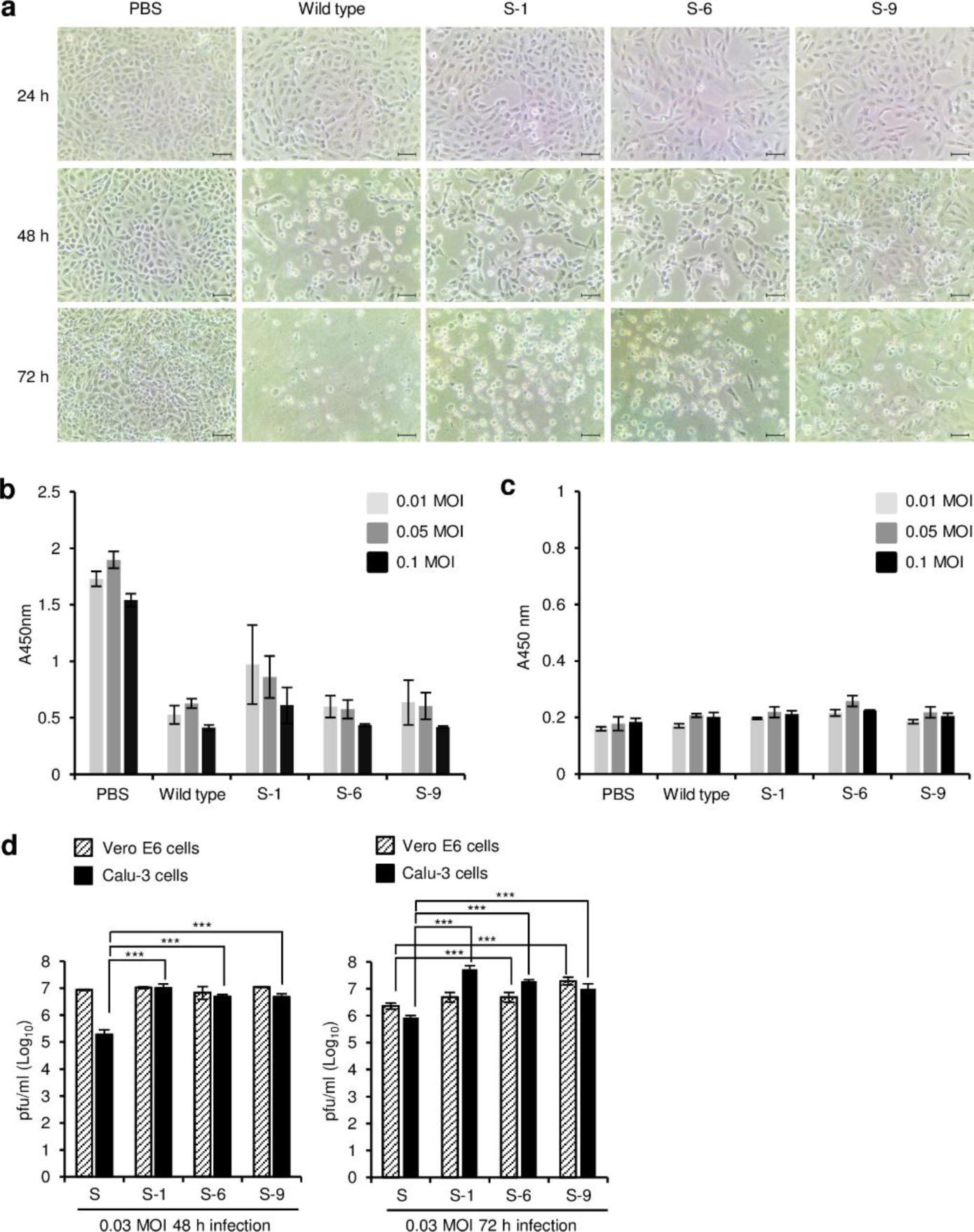
Differential replication of wild-type SARS-CoV-2 and its variants in Vero E6 and Calu-3 cells. **a,** Microscopic observation of SARS-CoV-2–infected Vero E6 cells. Vero E6 cells were infected with wild-type SARS-CoV-2 or its variants at 0.05 multiplicity of infection (MOI). At the indicated times after infection, the cells were observed with a bright light microscope. Scale bar, 50 μm. **b,c,** Cell viability assay. Vero E6 (B) and Calu-3 (C) cells were infected with wild-type SARS-CoV-2 or its variants (n = 3) at the indicated MOI. After 72 h, cell viability was determined by CCK-8 assay. Data are presented as mean ± s.d. **d,** Replication of SARS-CoV-2 in Vero E6 and Calu-3 cells. Vero E6 and Calu-3 cells (2 × 10^5^ cells/well on 24-well plates) were infected with wild-type SARS-CoV-2 (S) or its variants (S-1, S-6, S-9) in PBS at an MOI of 0.03 for 1 h (n = 3). At the indicated times after infection, supernatants were collected and the virus titers were determined by plaque assay. Data are presented as mean ± s.d. The *P* values were determined by two-sided unpaired t-test. \*\*\**p* < 0.001. Representative data are shown from at least 2 independent experiments (**a-d**).

### Properties of SARS-CoV-2 variants in the K18-hACE2 mouse model

To estimate the lethality of variants that arose during persistent infection in the mouse xenografts, we infected K18-hACE2 mice with various concentrations of the three clones isolated from the xenografts and determined the LD_50_ (Supplementary Data Fig. 3). The LD_50_ of clones S-1, S-6, and S-9 were 2.0 × 10^4^, 3.2 × 10^3^, and 1.3 × 10^4^, respectively, whereas the LD_50_ of the wild type was 1.3 × 10^2^. To further estimate the pathological properties of the clones, we infected K18-hACE2 mice with wild-type virus or one of the three clones and monitored them according to the experimental scheme shown in Fig. 5a. When the mice were infected with a lethal dose of wild-type SARS-CoV-2 (3 × 10^4^ pfu), loss of body weight was clearly detected, and all the mice died by 9 days after infection (Fig. 5b,c). By contrast, when the mice were infected with the same dose of one of the clones, the resulting weight loss was relatively mild, and some of the mice survived. When we measured virus titers 5 days after infection, the virus titers in turbinate were similar for the wild type and the clones (Fig. 5d), but the titers in the lungs were lower for the clones than for the wild type (Fig. 5e). At 14 dpi, the sera of surviving mice infected with each of the clones contained IgG reactive to the wild-type S protein RBD (Fig. 5f), suggesting that infection with the clones induced production of antibodies that were cross-reactive to the wild-type virus.

**Fig. 5.**
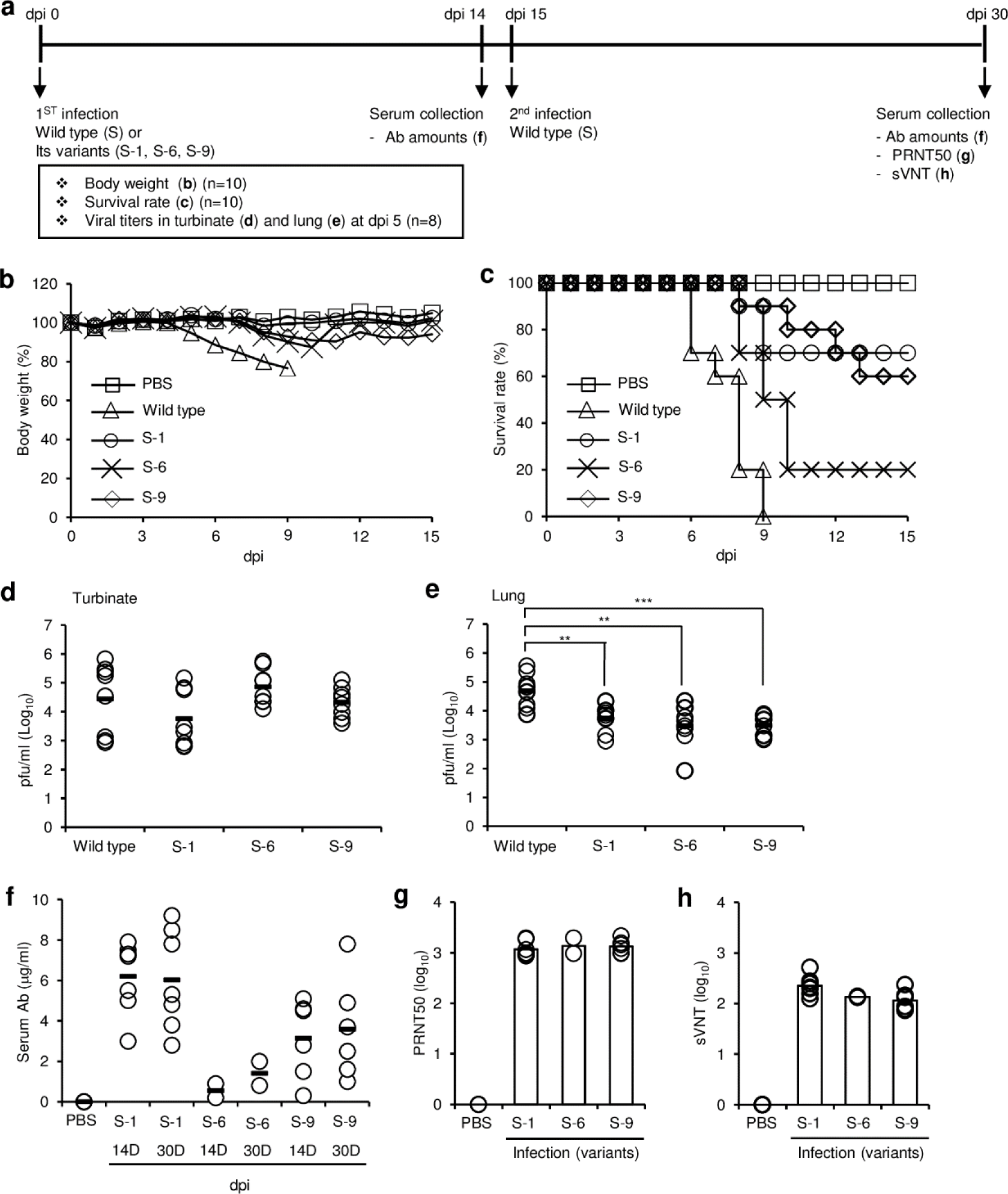
Effects of wild-type SARS-CoV-2 and its variants on hACE2 transgenic mice (B6.Cg-Tg(K18-ACE2)2Prlmn/J). **a,** Schematic diagram of the experiments. **b–e,** Hemizygous K18-hACE2 mice were intranasally infected with 3 × 10^4^ pfu/mouse of wild-type SARS-CoV-2 or its variants. Body weight (**b**) and survival (**c**) were measured for 15 days after intranasal infection (n = 10 per group). Data are presented as mean ± s.d. dpi, days post-infection. Turbinates (**d**) and lungs (**e**) were collected 5 days after intranasal infection (n = 8 per group). Viral loads in turbinate and lung homogenates were measured by plaque assay. Data are presented as mean ± s.d. The *P* values were determined by two-sided unpaired t-test. \*\**p* < 0.05, \*\*\**p* < 0.001. **f-h,** Hemizygous K18-hACE2 mice were intranasally infected with 3 × 10^4^ pfu/mouse of wild-type SARS-CoV-2 or its variants (n = 10 per group). Serum was collected from surviving mice 14 days (14D) after infection. Fifteen days after the first infection, the surviving mice were intranasally challenged with 3 × 10^4^ pfu/mouse of wild-type SARS-CoV-2. Serum was collected 15 days (30D) after the challenge. **f,** The amounts of SARS-CoV-2 wild-type S protein RBD-specific IgG in the serum were determined by ELISA. **g,** The PRNT_50_ of the serum collected at 15 days (30D) after the challenge was determined by plaque assay. **h,** The sVNT of the serum collected at 15 days (30D) after the challenge was determined against wild-type SARS-CoV-2. Data are presented as mean ± s.d (f-h).

To investigate whether prior infection with one of the clones could protect the mice against subsequent challenges with the wild-type virus, we challenged the surviving mice with a potentially lethal dose of wild-type virus and found that all the mice were still alive 15 days after the challenge. At 15 days after the challenge with wild-type virus, the amounts of serum antibodies were similar to or slightly higher than those before the challenge, suggesting that there was no boosting effect. However, PRNT_50_ and sVNT assays showed that the mice had substantial amounts of neutralizing antibodies against the wild-type virus in their serum (Fig. 5g,h). These results suggest that the lethality of the clones was lower than that of the original wild type, and infection with the clones provided protective immunity against subsequent challenge with the more lethal wild-type virus.

### Confirmation of SARS-CoV-2 persistent infection in the mouse xenograft model

To assess the variation of viral sequences that emerged among individual mice, we infected mouse xenografts (n = 6) with wild-type virus and confirmed persistent infection in the tumors based on virus titers and expression of viral proteins in the tumor tissues (Supplementary Data Fig. 4). We obtained virus samples from the tumor tissues at 3, 15, and 30 dpi and analyzed them by whole-genome sequencing. The viral sequences showed an overall similar pattern of mutations in the S protein with only minor differences compared with Fig. 2 (Fig. 6 and Supplementary Data Table 3). However, the mutation frequency at each site in ORF1ab, ORF3a, ORF7a, and ORF8 differed among the individual mice (Supplementary Data Table 2,3 and Supplementary Data Fig. 5,6). S protein mutations E484D and H655Y were found in all six mice with high frequency (E484D over 96%, H655Y over 88%), whereas the R685S mutation was not found in any mice, which is in line with the results in Fig. 2. The mutations T76I, T95I, N148K, and N185K were common and varied in frequency among the mice. Several additional mutations were found in the S protein [L5F, A27V, H146Q, N149T, D178N, Del(141-143LGV)], ORF1ab, ORF3a, ORF7a, and ORF8 at 30 dpi compared with Fig. 2. The S protein mutations found in these mice were previously found in several variants (Supplementary Data Table 1). Overall, we confirmed with multiple mice that diverse viral mutations, including some previously reported ones, occurred in the xenograft model, and some mutations were selected *in vivo*.

**Fig. 6.**
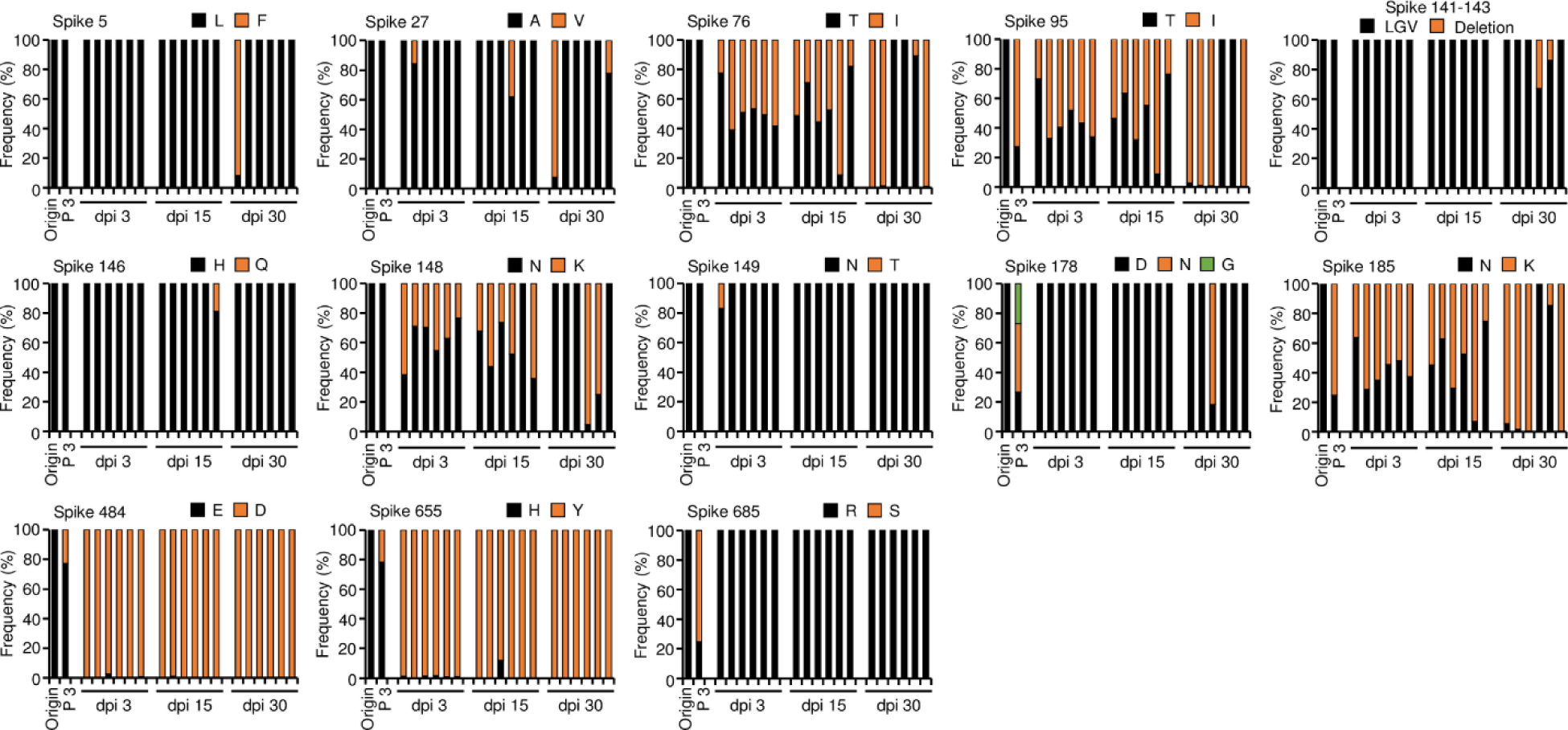
Frequencies and substitutions of amino acid sequences in wild-type SARS-CoV-2 S protein during long-term infection in lung tumor tissues. Calu-3 cells were subcutaneously implanted into the right flank of NRGA mice (n = 6 per group). After the tumor volume reached 100 mm^3^, the tumors were inoculated with 1 × 10^6^ pfu of wild-type SARS-CoV-2. Whole-genome sequences of the viruses in the tumors were analyzed at 3, 15, and 30 dpi. Each panel indicates the accumulation and substitutions of mutations at an amino acid position in SARS-CoV-2 S clade during long-term infection in lung tumor tissues. Origin: amino acid sequences of wild-type SARS-CoV-2 (Supplementary Table 1). Passage 3 (P 3): amino acid sequences analyzed in wild-type SARS-CoV-2 after amplification in three passages in Vero E6 cells. dpi, days post-infection.

## Discussion

RNA viruses have high mutation rates because of the low fidelity of RNA-dependent RNA polymerase. These viruses also encounter host genetic variation and diverse cellular microenvironments during outbreaks of infection. During the last 3 years, continuous waves of several SARS-CoV-2 variants have spread around the world. Several hypotheses were suggested to explain this active emergence of SARS-CoV-2 variants, including persistent infection in immunocompromised patients^23^, simultaneous infection and recombination of variants in unvaccinated or immunocompromised individuals^17^, and transmission between humans and animals resulting in accelerated evolution^24^. We are interested in the first hypothesis, as persistent infection of SARS-CoV-2 and the occurrence of its variants in patients with COVID-19 were previously reported by many investigators^11–14^.

There are animal models to recapitulate symptoms and immune responses after SARS-CoV-2 infection, including human ACE2-transgenic mice, hamsters, ferrets, and non-human primates^25^. These models are useful to study the pathogenesis, virulence, and immune escape of variants of concern and the efficacy of vaccines and therapeutics against these variants. However, there is an unmet need for an animal model to recapitulate persistent infection in which variants of SARS-CoV-2 can emerge after long-term replication *in vivo*^26^. Therefore, we established a novel persistent infection model using xenograft tumor tissues derived from Calu-3 cells.

The first evidence suggesting that our model recapitulates persistent SARS-CoV-2 infection *in vivo* is the continuous production of viruses in the tumor tissues. When we established xenografts with Calu-3 cells and infected them with SARS-CoV-2, the mice showed no prominent side effects of virus infection, although high titers of virus were present in the tumor tissues for 30 days, which was the endpoint of our experiment. We did not measure virus titers after 30 dpi; however, persistent infection in the mouse xenografts may continue much longer, as it does in human patients with SARS-CoV-2 infection^11–14^. We believe the selection of appropriate cells that do not suffer cytotoxic effects due to SARS-CoV-2 infection was the most important factor in our model. As we reported previously, Calu-3 cells replicated continuously without apoptosis for 3 days after SARS-CoV-2 infection, in contrast to Vero cells that displayed cytotoxic effects after infection^15^. Calu-3 cells are cancer cells derived from the human lung, a primary target organ of SARS-CoV-2, but SARS-CoV-2 has also been found in other organs in humans including the gastrointestinal tract, liver, and brain^27^. Furthermore, extensive autopsy data suggest that SARS-CoV-2 causes systemic infection in humans with a wide range of severity and can persist in various organs for several months^28^. As human cell lines such as Caco-2 colorectal adenocarcinoma cells, Huh-7 liver cancer cells, and U251 glioblastoma cells are susceptible to SARS-CoV-2 without cytopathic effects^27^, we speculate that other cell lines can be used in xenograft infection models to provide more insights into persistent infection in different human tissues.

Viruses in the xenografts randomly displayed many mutations at several residues, similar to virus pools in cell culture. Some critical mutations reached high frequency in the xenograft infections, whereas others were displaced by the original sequences. For example, all viral clones isolated from the xenografts showed the S protein mutation H655Y and no mutation at residue S685, and this phenomenon was not observed in Vero E6 cell cultures. It can be understood that passage in cell culture provides a kind of artificial amplification of a small fraction of viral genotypes, but propagation in a xenograft model can show virus evolution *in situ*. Therefore, we conclude that there were continuous changes in the virus pool in the mouse xenografts, with some unique changes that differed from those that occurred during *in vitro* propagation.

Previously, mutations involved in SARS-CoV-2 entry into cells were evolved by serial passage in human cell lines without ACE2 or cellular proteases^22, 29^. Furthermore, several mutations reported in natural variants were recapitulated by serial passaging in several human cell lines, with some differences between colon epithelial cells and lung epithelial cells^30^. These results suggest the specific host cell environment can affect the selection of mutations. We believe that our *in vivo* persistent infection model recapitulates natural infection better than cell culture systems; however, the important key mutation D614G was not found in our study, which was the same as in a previous investigation^35^. Therefore, we speculate that the probability of this mutation is very low, but it was somehow selected in humans. It is also possible that our model has a limitation, as the xenografts are in a mouse host.

We analyzed a pool of viruses isolated from xenograft tumor tissues at 30 dpi by whole-genome sequencing. The gene for the S protein covers only about 12.8% of the SARS-CoV-2 genome; however, 60% of the mutations that differentiate the Omicron variant from the original Wuhan strain are in the S protein^31^. This trend was also found in our results. The sporadic mutations in the viral genomes isolated from multiple mice were mostly found in the S protein, especially the S1 region, and most of these mutations were previously reported in natural variants or variants obtained by cell culture. The functional effects of the identified mutations on virus properties are largely unknown, except for E484D and H655Y. The E484D mutation was implicated in ACE2-independent entry of SARS-CoV-2 into cells^20^. Furthermore, mutations at this site (E484A, E484D, E484G, and E484K) contribute to escape from neutralization by antibodies from convalescent plasma^32^. The S protein H655Y mutation is expected to enhance the endosomal entry pathway and contribute to the low pathogenicity of the Omicron variant^33^. Mutations in the S protein are important in the infection, pathogenicity, and immune escape of SARS-CoV-2 variants^39–41^. The D614G mutation induces higher infectious titers by increasing the stability of the S protein and promoting higher incorporation of S protein into virions without changing the affinity of the S protein for ACE2^34, 35^. The RBD (amino acids 333–527) of the S protein is the target of 90% of neutralizing antibodies in the sera of patients infected with SARS-CoV-2^36^. The mutation N439K, located in the receptor binding motif (amino acids 438–506) of the S protein, increases the binding affinity of the S protein to ACE2 and reduces the neutralizing activity of antibodies in the sera of patients with SARS-CoV-2 infection. Therefore, it is necessary to further investigate the functional effects of the major mutations that we found in our persistent infection model. In addition, further investigation of the isolated clones with multiple mutations will provide insights into the combined effects of the mutations. Such efforts may help to predict the properties of emerging variants once a genomic sequence is obtained without any further knowledge of the variants. Although we focused on S protein mutations, we also found several mutations in the N protein, ORF1ab, ORF3a, ORF6, ORF7a, and ORF8. Considering that mutations modulating protein levels of innate immunity antagonists such as the N protein, ORF9b, and ORF6 enhance viral escape from host innate immunity^37^, our results might provide clues about additional genes and mechanisms contributing to the properties of variants.

To obtain more information about the variants obtained in our persistent infection model, we analyzed individual viral clones by whole-genome sequencing, plaque assays, and *in vivo* challenge experiments. The whole-genome sequencing revealed that viral plaques were not clones composed of a single unique genome but rather a mixed population of viruses from the same origin, supporting the occurrence of continuous mutation in SARS-CoV-2. Therefore, the term “clone” that we used in this study is not exactly right, and “quasi-clone” might be more accurate. However, we used the term to emphasize that we isolated viruses from single isolated plaques. The viral clones had different combinations of mutations, so the combined effects of the mutations can be measured using these viruses. The viral clones obtained from xenograft tissues showed higher replication in Calu-3 cells and decreased lethality in mouse challenge experiments compared with the original wild-type SARS-CoV-2. These results support general speculation that SARS-CoV-2 variants acquire mutations that promote increased replication and decreased lethality as they adapt to their human host. However, the occurrence of frequent sporadic mutations and the accumulation of some critical mutations during persistent infection may result in the emergence of variants of concern. In addition, other factors such as recombination of variants and cross-species transmission may accelerate this process^21, 29^. Taken together, our results suggest that Calu-3 xenograft mice can be used as a model to study persistent infection of SARS-CoV-2 and its variants.

## Methods

### Cell culture and virus

Human airway epithelial Calu-3 cells (cat. no. 30055), adenocarcinomic human alveolar basal epithelial A549 cells (cat. no. 10185), and African green monkey kidney Vero E6 cells (cat. no. 21587) were obtained from the Korean Cell Line Bank. The cells were maintained in Dulbecco’s modified Eagle’s medium (DMEM; Thermo Fisher Scientific) containing 10% fetal bovine serum (FBS; Thermo Fisher Scientific), 25 mM HEPES, 100 U/mL penicillin, and 100 μg/mL streptomycin at 37°C in a 5% CO_2_ incubator. Wild-type SARS-CoV-2 (hCoV-19/Korea/KCDC03/2020, lineage A, NCCP43326, GenBank accession no. MW466791.1) was provided by the National Culture Collection for Pathogens of Korea.

### Virus amplification and determination of virus titers by plaque assay

SARS-CoV-2 amplification in Vero E6 cells was performed as described previously^15, 38^. Vero E6 cells (7 × 10^5^ cells/well) were cultured overnight on six-well plates (Corning) in DMEM containing 10% FBS. The cells were then infected with SARS-CoV-2 at a multiplicity of infection (MOI) of 0.01 for 1 h at 37°C with shaking every 10 min in a CO_2_ incubator. DMEM containing 2% FBS was then added, the cells were cultured for 72 h, and the culture supernatants were collected by centrifugation. Virus titers in the culture supernatants and homogenates of tumor tissues were quantified by plaque assay on Vero E6 cells as described previously^39, 40^. Virus stocks (1 × 10^7^ pfu/mL) were stored at −70°C. SARS-CoV-2 amplification and cell culture procedures were performed in biosafety level 3 (BSL-3) conditions at the Research Institute of Medical-Bio Convergence of Hallym University with approval of the Institutional Biosafety Committee (IBC) of Hallym University (Hallym2020-12, Hallym2021-04, Hallym2022-03).

### Cell viability assay

Vero E6 cells (5 × 10^3^ cells/well) or Calu-3 cells (5 × 10^3^ cells/well) were cultured on 96-well plates for 18 h. The cells were then washed with PBS and infected with SARS-CoV-2 in PBS at an MOI of 0.01, 0.05, or 0.1 for 1 h in a 5% CO_2_ incubator at 37° with shaking every 20 min. The cells were then washed with PBS, and DMEM containing 2% FBS was added to each well. After 72 h, the viability of SARS-CoV-2–infected cells was assessed by cell-counting kit-8 (CCK-8) assay (Dojindo Laboratories, CK04). After adding CCK-8 reagent, cell viability was determined by measuring the absorbance at 450 nm following the manufacturer’s instructions.

### Animals

For the mouse xenograft model, four-week-old female NOD/ShiLtJ-Rag2*^em1AMC^*Il2rg*^em1AMC^* (NRGA) mice were purchased from JA BIO, Inc. For the virus challenge experiments, seven-week-old male hACE2 transgenic mice (B6.Cg-Tg(K18-ACE2)2Prlmn/J) were purchased from The Jackson Laboratory (stock. no. 034860). All mice were maintained under specific pathogen-free conditions in a controlled environment (20–25°, 40–45% humidity, 12-h light/dark cycle, *ad libitum* access to food and water) in the Experimental Animal Center of Hallym University. Humane endpoints were planned whereby the mice would be anesthetized by intraperitoneal injection of 0.2 mL avertin (Sigma-Aldrich) and euthanized by cervical dislocation in accordance with the approved Institutional Animal Care and Use Committee (IACUC) protocol if they lost 30% of adult body weight, reached a tumor volume of 1,000 mm^3^, or exhibited evidence of debilitation, pain, or distress such as hunched posture, rough haircoat, reduced food consumption, emaciation, inactivity, difficulty ambulating, or respiratory problems. No live mice reached the humane endpoint criteria during the experiments. All animal procedures were approved by the IACUC of Hallym University (Hallym2020-26, Hallym2021-73). Animal experiments involving SARS-CoV-2 infection were performed in animal biosafety level 3 (ABSL-3) conditions in the Research Institute of Medical-Bio Convergence of Hallym University in accordance with the recommendation of the IBC of Hallym University (Hallym2020-12, Hallym2021-04, Hallym2022-03).

### Mouse xenograft model and virus infection experiments

Calu-3 cells or A549 cells (2 × 10^6^ cells/mouse) in 50% Matrigel (Corning, PBS/Matrigel, 1:1 *v*/*v*) were subcutaneously inoculated into the right flank of four-week-old female NRGA mice. After tumor volumes reached an average of 100 mm^3^, the mice (n = 3/group or n = 6/group) were intratumorally infected with 1 × 10^6^ pfu of wild-type SARS-CoV-2. Then, the mice were monitored daily for body weight, clinical signs, and survival. Tumor diameters were measured with calipers. The tumor volumes were calculated as width^2^ × length/2 as described previously^41^. The mice were sacrificed 3, 15, or 30 dpi, and the tumors were surgically excised and weighed. To measure the virus distribution in the infected mice, tumors, lungs, brains, and blood were collected at 30 dpi. Each organ was homogenized using Tissue Lyser II (Qiagen), and the virus titer of each organ was determined by plaque assay and quantitative real-time reverse-transcription PCR (qRT-PCR).

### Western blot

SARS-CoV-2–infected tumors were homogenized and lysed in cell lysis buffer (10 mM HEPES, 150 mM NaCl, 5 mM EDTA, 100 mM NaF, 2 mM Na_3_VO_4_, protease inhibitor cocktail, and 1% NP-40). Cell lysates were then collected by centrifugation. Equal amounts of protein were loaded and resolved on a 4–12% Bis-Tris gradient gel (Thermo Fisher Scientific) and analyzed by western blot with anti-SARS-CoV-2 N protein monoclonal antibody (0.5 μg/mL, anti-SARS-CoV-2 N mAb, clone 1G10C4 mAb)^42^ and anti-β-actin antibody (Sigma-Aldrich, cat. no. A5316, RRID: AB_476743 (1:5000)). A horseradish peroxidase (HRP)-conjugated donkey anti-mouse IgG (H+L) polyclonal antibody (Jackson ImmunoResearch Laboratories, Inc., cat. no. 715-035-150, RRID:AB_2340771, (1:5000)) was used as secondary antibody. Immunoreactive bands were detected using an enhanced chemiluminescence (ECL) reagent (Thermo Fisher Scientific)

### qRT-PCR

SARS-CoV-2–infected tumors were homogenized, and the supernatants were collected. Total RNAs were extracted from the supernatants using TRIzol Reagent (Invitrogen) and a PureLink™ RNA Mini Kit (Invitrogen). Then, cDNA (50 μL) was synthesized using a Reverse Transcription System Kit (Promega). Expression of the RNA-dependent RNA polymerase (*RdRP*) gene of SARS-CoV-2 (wild type) was quantified using the following primers as previously described^39^ : forward primer, 5′-GTGAAATGGTCATGTGTGGCGG-3’; reverse primer, 5’-CAAATGTTAAAAACACTATTAGCATA-3’; TaqMan probe, 5’-FAM-CAGGTGGAACCTCATCAGGAGATGC-TAMRA-3’. Primers and the probe were synthesized by Genotech. 10 µL of GoTaq Probe qPCR Master Mix (Promega, A6101) was added to 10 µL of the reaction mixture containing 125 nM each of the forward and reverse primers, 250 nM probe, and 1 μL of cDNA. After denaturation at 95° for 5 min, 45 cycles at 95° for 15 sec and 60° for 1 min were performed in the Rotor-Gene Q real-time PCR cycler (Qiagen). The numbers of copies of the *RdRP* gene in the samples were calculated using an *RdRP* cDNA standard curve.

### Hematoxylin and eosin (H&E) staining

Tissue sections were prepared for histopathologic examination as previously described^43^. In brief, tumor tissues were removed at 3, 15, 30 days after infection and fixed in a 4% paraformaldehyde. The tissues were embedded in paraffin and sectioned into 5 µm slices. After the tissue sections were allowed to dry on glass slides overnight at 40°C, the slides were melted, deparaffinized, and rehydrated. The rehydrated slides were stained with Gill’s Hematoxylin V (Muto Pure Chemicals), and counterstained with Eosin Y solution (Abcam). The slides were then dehydrated with ethanol and then the slides were treated with xylene and mounted with a 6:4 solution of Malinol (Muto Pure Chemicals) and xylene. The H&E-stained sections were observed using an Eclipse E200 microscope (Nikon).

### Immunohistochemistry

Paraffin-embedded tissue sections were deparaffinized and rehydrated. The rehydrated slides were treated with 3% H_2_O_2_ and boiled for antigen retrieval in citrate buffer (pH 6.0: ScyTek Laboratories). Each slide was then blocked with avidin blocker, biotin blocker (Vector Laboratories), and normal horse serum (Vector Laboratories). Then, each slide was incubated with biotinylated anti-SARS-CoV-2 N monoclonal antibody (1G10C4 mAb, 1 μg/slide)^44^, rabbit anti-SARS-CoV-2 S polyclonal antibody (Sino Biological, cat. no. 40591-T62, RRID:AB_2893171, 1 μg/slide), or rabbit anti-ACE2 polyclonal antibody (MyBioSource, cat. no. MBS4750512, (1:200)). The slides stained with rabbit anti-SARS-CoV-2 S polyclonal antibody or rabbit anti-ACE2 antibody were further incubated with biotinylated horse anti-rabbit antibody (Vector Laboratories, cat. no. BP 1100). All slides were then incubated with HRP-conjugated streptavidin (Vector Laboratories, cat. no. PK-6100, RRID: AB_2336819, (1:200)). The enzyme-substrate reaction was developed using 3, 3’-diaminobenzidine (DAB, Thermo Fisher Scientific). The slides were counterstained with Gill’s Hematoxylin V. The H&E-stained sections were observed using an Eclipse E200 microscope.

### Viral RNA extraction from tumor tissues and whole-genome analysis

Total RNA samples were extracted from the supernatants of virus-infected tumor homogenates and sent to Macrogen, Inc. for cDNA library construction and whole-genome sequencing. The libraries were quantified using Kapa Library Quantification kits for Illumina Sequencing platforms (Kapa Biosystems). Sequencing was performed using an Illumina NovaSeq system (Illumina, Inc.). The sequencing data were filtered with FastQC (v0.11.8) (https://www.bioinformatics.babraham.ac.uk/projects/fastqc/) and Trimmomatic (v0.38)^45^ to check the read quality and reduce biases for analysis. Filtered data were mapped to reference sequences using BWA (v0.7.17)^46^ with the MEM algorithm and modified with Sambamba (v0.6.7)^47^. SAMtools (v.1.6)^48^ and BCFtools (v.1.6) (http://samtools.github.io/bcftools/call-m.pdf) were used for genome coverage determination, mapping ratio calculation, and variant calling. At this step, single-nucleotide polymorphisms and short insertion or deletion candidates with Phred score over 30 (base call accuracy of 99.9%) were captured and annotated with SnpEff (v.4.3t)^49^ to predict the effects of genetic variants. The sequences of wild-type SARS-CoV-2 (hCoV-19/Korea/KCDC03/2020, lineage A, NCCP43326/2020, GenBank accession no. MW466791.1.) were used as a reference. The accession numbers for the SARS-CoV-2 wild-type proteins are listed in Supplementary Table 1.

### Analysis of the S protein sequences of the variants by reverse-transcription PCR

SARS-CoV-2–infected tumor tissues were homogenized, and the supernatants were collected. Total RNAs were extracted from the supernatants of the homogenates, and cDNA was generated. For cloning of the S protein N-terminal region including the RBD (2726 bp, from −46 to 2680), standard PCR was performed for 25 cycles using AccuPrime^TM^ Taq DNA polymerase (Invitrogen) and the following primer sets with Not I and Kpn I restriction sites included at the 5′ and 3′ ends, respectively: CoV2-S-5’ primer, 5’-TATA<u>GCGGCCGC</u>CAGAGTTGTTATTTCTAGTGATGTTC-3’; CoV2-S-mid-3’ primer, 5’-TATA<u>GGTACC</u>ATGCAGCACCTGCACCAAAG-3’. The PCR products were inserted into a modified pcDNA 3.4 expression vector (Thermo Fisher Scientific), and the nucleotide sequences were verified by DNA sequencing using the following primers: S1-5’p forward primer, 5’-CTGGTGATTCTTCTTCAGGT-3’; S2-5’p forward primer, 5’-GAACTTCTACATGCACCAGC-3’; S0-3’p reverse primer, 5’-GTGCACAGTCTACAGCATCT-3’; pcDNA3.4 reverse primer, 5’-CAACATAGTTAAGAATACCAGTC-3’.

### Isolation of viral plaques and sequence analysis of viral clones

SARS-CoV-2–infected tumors were homogenized, and the supernatants were collected. For viral plaque isolation, serially diluted supernatants were supplemented onto Vero E6 cells (7 × 10^5^ cells/well) on six-well plates. After 1 h of infection, 3 mL/well DMEM/F12 medium (Thermo Fisher Scientific) containing 2% Oxoid agar was added. After incubation at 37°C for 3 days, five plaques from each homogenate of three tumors were collected, and each clone was amplified in Vero E6 cells. The S protein sequences of the clones were analyzed by reverse-transcription PCR and DNA sequencing, and three variants (S-1, S-6, and S-9) were selected and analyzed further by whole-genome sequencing. The virus stocks of the variants were stored at −70°C.

### Virus challenge experiments

Eight-week-old male hemizygous K18-hACE2 mice were anesthetized by exposure to 1–2% isoflurane (Hana Pharm. Co.) and then intranasally inoculated with wild-type SARS-CoV-2 or variant. To investigate the LD_50_ for each virus, mice (n = 5/group) were intranasally inoculated with serial dilutions of the viruses and monitored daily for clinical signs, body weight, and survival for up to 15 days. The LD_50_ was defined as the dilution of the virus that produced lethality in 50% of the mice. LD_50_ titers were calculated by the method of Reed and Muench^50^. At 15 dpi with 3 × 10^4^ pfu of SARS-CoV-2 wild type or variant (n = 10/group), live mice were intranasally challenged with 3 × 10^4^ pfu of wild-type SARS-CoV-2 and then monitored for another 15 days for clinical signs and body weight^51^. To investigate virus-specific antibody production and neutralizing antibody titers, blood samples were collected by retro-orbital bleeding 14 days after the initial intranasal inoculation and by cardiac puncture 15 days after the subsequent intranasal challenge. Serum samples were prepared and stored at −80°C. To examine the viral loads after inoculation with SARS-CoV-2 wild type or variant, mice (n = 8/group) were sacrificed 5 days after intranasal inoculation, and turbinates and lungs were removed for analysis. The turbinates and lungs were weighed and homogenized, and virus titers were estimated by plaque assay.

### RBD-specific Ig ELISA

Ninety-six–well immunoplates (Nunc^TM^) were coated with recombinant RBD of SARS-CoV-2 wild-type S protein (1 μg/mL, Catalog No. SPD-C82E9, Acrobiosystems) in carbonate buffer (pH 9.6) and incubated overnight at 4°C. The coated plates were blocked with PBST containing 3% BSA and then washed three times with PBST. Twofold serially diluted sera were then transferred to each well and incubated for 2 h at room temperature. After incubation, the plates were washed and incubated with HRP-conjugated goat anti-mouse IgG antibody (Southern Biotechnology Associates, cat. no.1030-05, RRID: AB_2619742 (1:200)). After incubation for 1 h at room temperature, the plates were washed five times with PBST, and the colorimetric reaction was developed using the substrate 3,3’,5,5’-tetramethylbenzidine (TMB; Kirkegaard and Perry Laboratories). After optimal color development, the reaction was stopped using a stop solution (Sera Care Life Sciences, Inc), and the absorbance was measured at 450 nm with a Spectra Max 250 microplate reader (Molecular Devices)

### Plaque reduction neutralization test (PRNT)

Vero E6 cells (7 × 10^5^ cells/well) were cultured on six-well plates for 18 h. SARS-CoV-2 wild type was pre-incubated with twofold serially diluted mouse serum for 1 h at 37°C. After the Vero E6 cells were washed with PBS, the virus-serum mixture was added. After the cells were infected for 1 h with shaking every 10 min, the supernatant was removed and 3 ml of DMEM/F12 medium (Thermo Fisher Scientific) containing 2% Oxoid agar was added. The plates were cultured for 72 h and then stained with 0.1% crystal violet to visualize plaque formation. Antibody titers were calculated using the Spearman-Kärber method^52^ and percentage of inhibition ≥50% (PRNT_50_) is considered a positive cutoff for seroconversion against SARS-CoV-2.

### Surrogate SARS-CoV-2 virus neutralization test (sVNT)

Recombinant SARS-CoV-2 wild-type S protein RBD-HRP fusion protein (RBD-HRP protein, cat. no. Z03594) and hACE2 protein (cat. no. Z03516) were purchased from GenScript Korea Ltd. The sVNT was performed as previously described^53^. Briefly, hACE2 protein (1 µg/mL) was coated on a 96-well plate and then blocked with PBS containing 1% BSA. Then, 50 µ L RBD-HRP protein (2 µg/mL) in PBST and 50 µ L twofold serially diluted serum were mixed and incubated for 30 min. One hundred microliters of the mixture were then added to the hACE2-coated 96-well plate and incubated for 15 min. The plate was then washed with PBST, TMB peroxidase substrate was added, and absorbance at 450 nm was measured. The absorbance values were normalized with the values of PBS (negative control) and RBD-HRP protein alone (positive control) in hACE2-coated wells^54^. The surrogate virus neutralization titer (sVNT_50_) of the serum was defined as the reciprocal value of the sample dilution that showed a 50% reduction of signal at 450 nm.

### Statistical analysis

Results are shown as the mean ± standard deviation. Differences between the samples were analyzed using a two-sided unpaired Student’s t-test (Instat; GraphPad Inc.). *p*-values<0.05 were considered statistically significant.

### Data availability

All data needed to evaluate the conclusions in this manuscript have been included.

### Code availability

The data form whole genome sequencing was analyzed using publicly available tools. After viral genome samples were sequenced using Illumina NovaSeq platform, FastQC (v0.11.8) was used to check the read quality and Trimmomatic (v0.38) was used to remove adapter sequences and low-quality reads in order to reduce biases for analysis. Filtered data were mapped to reference using BWA (v0.7.17) with MEM algorithm. After the mapping process, duplicated reads were removed with Sambamba (v0.6.7). Genome coverage, mapping ratio calculation and variant calling were performed using SAMtools (v.1.6) and BCFtools (v.1.6). At this step, SNPs and short indels candidates with Phred score over 30 (base call accuracy of 99.9%) were captured and annotated with SnpEff (v.4.3t) in order to predict the effects of genetic variants.

## Acknowledgements

We thank the National Culture Collection for Pathogens (Osong, Korea) for supplying SARS-CoV-2 wild type. This research was supported by a grant from the National Research Foundation (grant number: NRF-2022M3A9I2082292) funded by the Ministry of Science and ICT in the Republic of Korea and by a grant of the Korea Health Technology R&D Project (grant number: HV22C0062) through the Korea Health Industry Development Institute (KHIDI), funded by the Ministry of Health & Welfare in the Republic of Korea.

## Author information

### Contributions

M.S.P., Y.L. and H.J.K. conceived the study. D.K., J.K., M.S.P., Y.L. and H.J.K. designed experiments. D.K., J.K., M.K., K.B., B.M.K. and S.K. performed experiments in BSL-3 and ABSL-3 facility including cell culture, viral cultivation and animal experiments. D.K. and J.K. carried out xenograft mouse model experiments. D.K., J.K. and M.K. carried out virus challenge experiment. J.K. and M.K. performed PRNT and sVNT experiments. M.K., K.B., B.M.K. and S.K. contributed to sample processing for DNA sequencing. D.K., J.K., S.M. and H.P performed viral RNA extraction, analyzed whole-genome sequences data and prepared figures. J.K. and S.P. performed qRT-PCR. K.B., B.M.K. and S.K. carried out H&E staining and immunohistochemistry. D.K., S.M., Y.L and H.J.K wrote the manuscript. All authors contributed to editing of the manuscript and approved the final manuscript.

---

**Supplementary Data Fig. 1.**
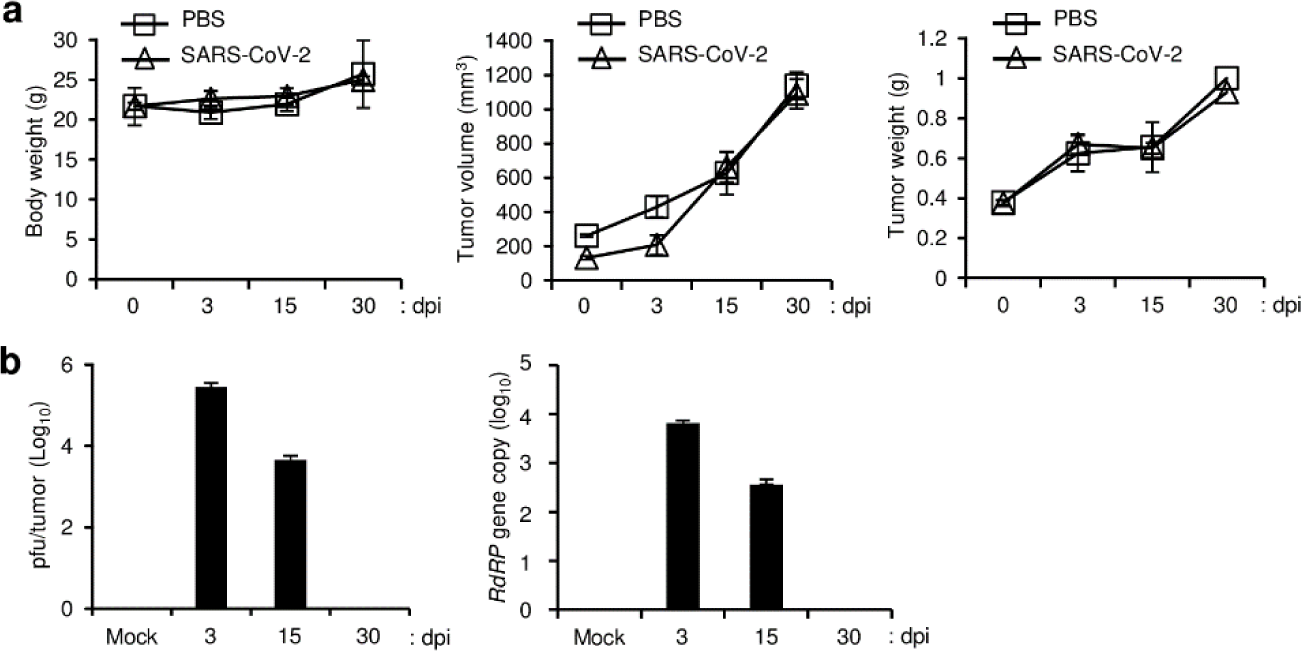
SARS-CoV-2 infection in an A549 cell-derived lung tumor xenograft mouse model. A549 cells were subcutaneously implanted into the right flank of NRGA mice (n = 3 per group). After the tumor volume reached 100 mm^3^, 1 × 10^6^ pfu of wild-type SARS-CoV-2 was inoculated into the tumors, and virus titers in the tumors were analyzed at 3, 15, and 30 dpi. **a,** Body weight of each infection group (left panel). Tumor volume of each infection group (middle panel). Tumor weight of each infection group (right panel). Data are presented as mean ± s.d. **b,** After SARS-CoV-2 infection, tumor tissues were collected at 3, 15, and 30 dpi. Viral titers in the supernatants of tumor homogenates were determined by plaque assay (left panel) and qRT-PCR analysis for the SARS-CoV-2 *RdRP* gene (right panel). Data are presented as mean ± s.d. dpi, days post-infection.

**Supplementary Data Fig. 2.**
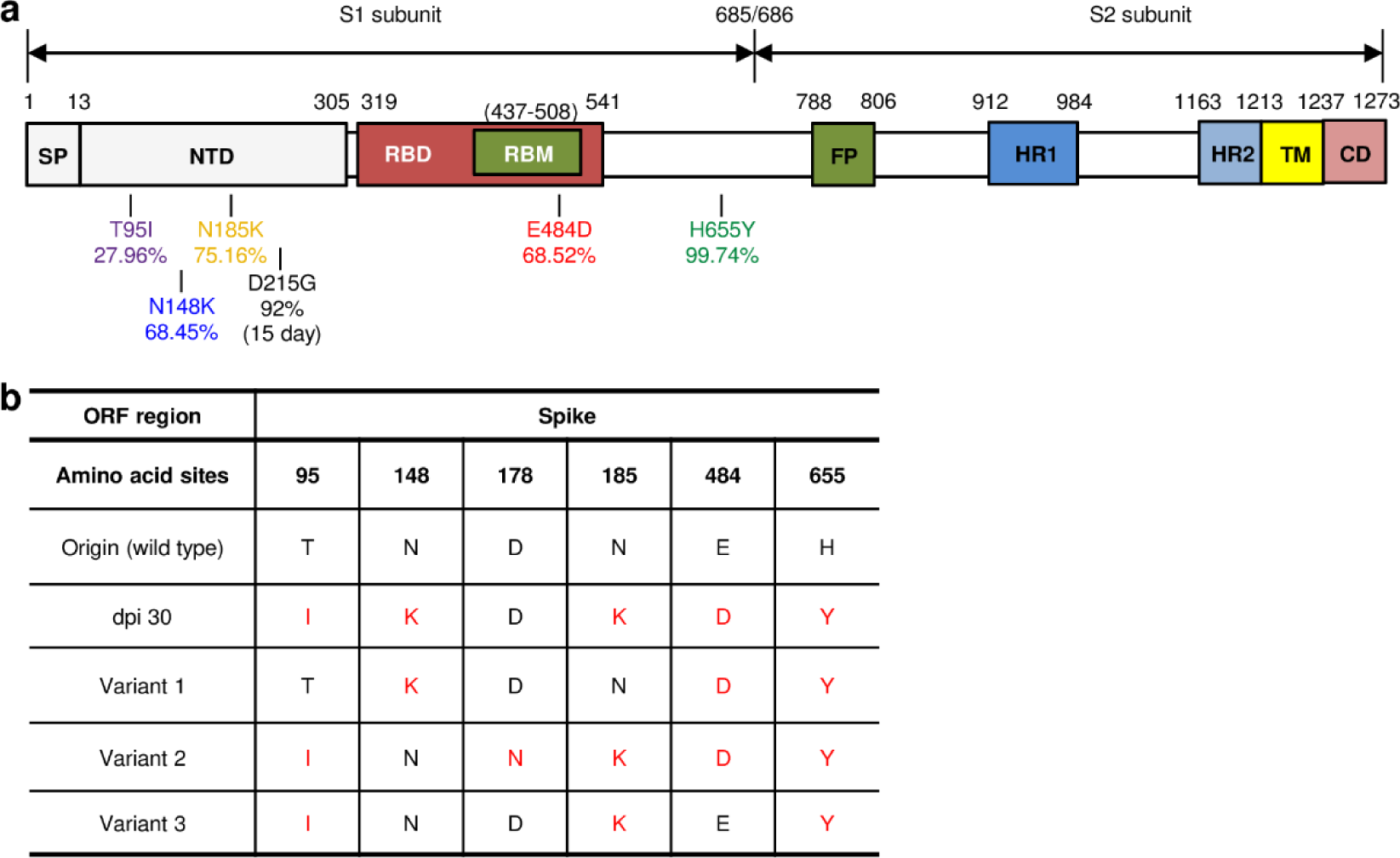
Analysis of amino acid mutations in the S protein of wild-type SARS-CoV-2 and its variants. **a,** Schematic diagram of amino acid mutations in the S protein of wild-type SARS-CoV-2. The mutations were analyzed by whole-genome sequencing of SARS-CoV-2 virus isolated from tumors at 30 dpi as shown in Figure 2. **b,** Analysis of S protein mutations generated in lung tumors. cDNA was synthesized from the pool of viruses in the supernatants of SARS-CoV wild-type-infected tumor homogenates obtained at 30 dpi. The sequences of the S1 subunit region of the S protein were cloned and analyzed by DNA sequencing. Origin (wild type): amino acid sequences of wild-type SARS-CoV-2 (Supplementary Table 1). Variant 1, Variant 2, and Variant 3: three types of variants containing amino acid mutations of the S1 subunit region of the S protein. dpi, days post-infection.

**Supplementary Data Fig. 3.**
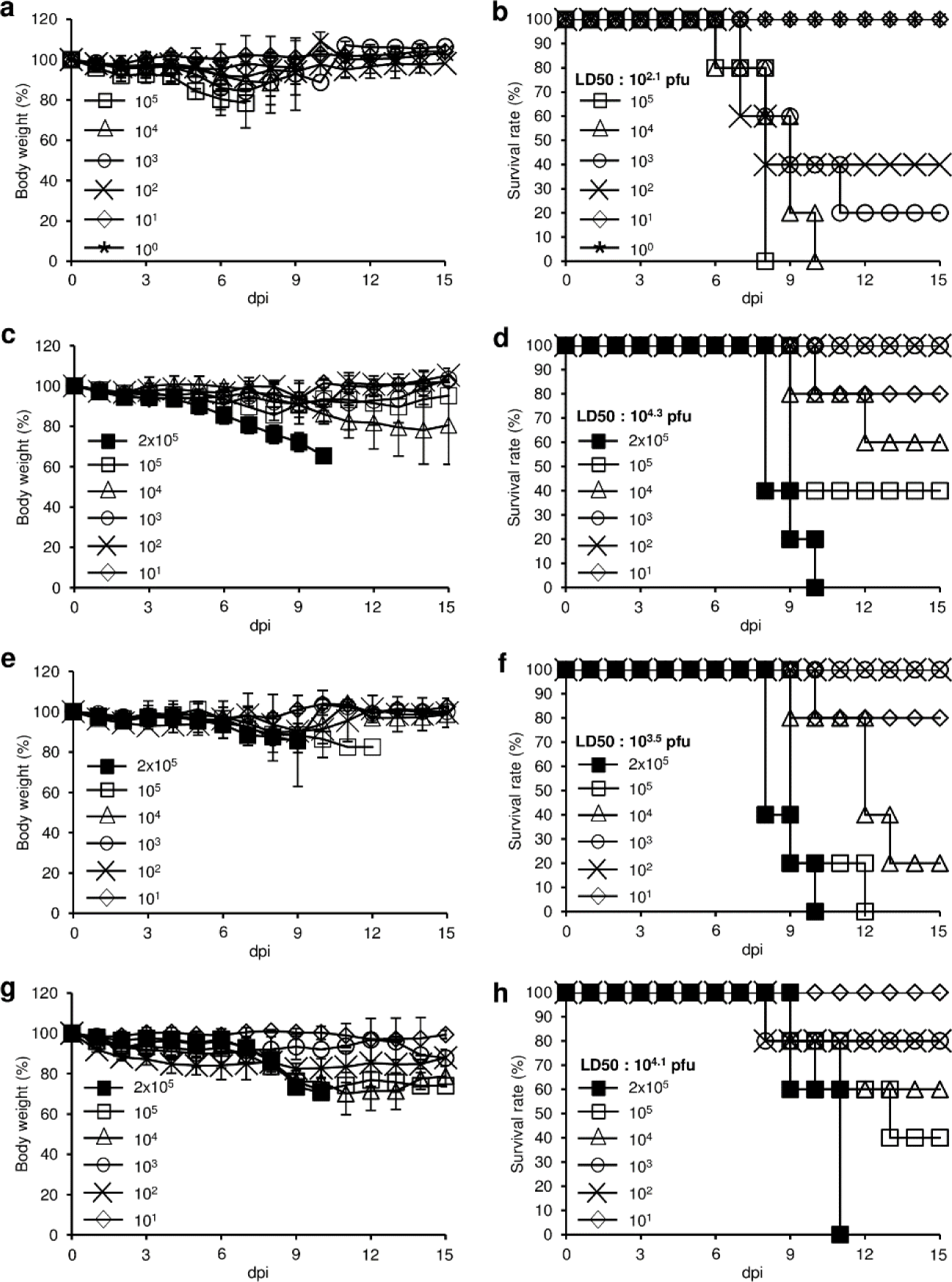
Determination of LD_50_ for each virus in K18-hACE2 mice. Eight-week-old male hemizygous K18-hACE2 mice (n = 5 per group) were intranasally inoculated with the indicated titers of wild-type SARS-CoV-2 or its variants. The mice were monitored daily for clinical signs and measured for body weight (a,c,e,g) and survival (b,d,f,h) for up to 15 days. **a,b,** wild-type SARS-CoV-2. **c,d,** S-1 variant. **e,f,** S-6 variant. **g,h,** S-9 variant. Data are presented as mean ± s.d. dpi, days post-infection.

**Supplementary Data Fig. 4.**
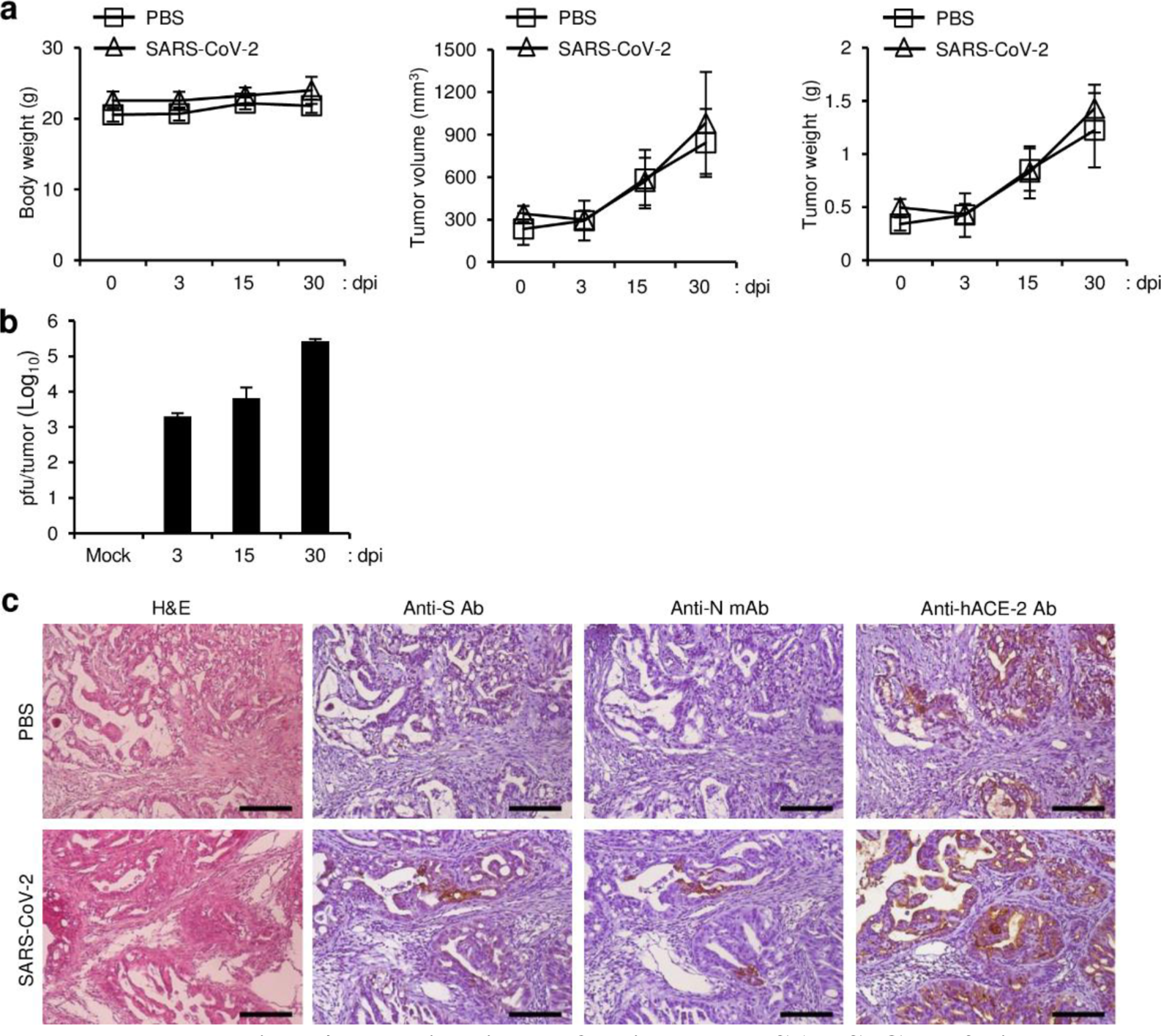
Replication of wild-type SARS-CoV-2 in the lung tumor xenograft mouse model. Calu-3 cells were subcutaneously implanted into the right flank of NRGA mice (n = 6 per group). After the tumor volume reached 100 mm^3^, the tumors were inoculated with 1 × 10^6^ pfu of wild-type SARS-CoV-2. Virus titers in the tumors were analyzed at 3, 15, and 30 dpi. **a,** Body weight in each infection group (left panel). Tumor volume in each infection group (middle panel). Tumor weight in each infection group (right panel). Data are presented as mean ± s.d. **b,** Replication of SARS-CoV-2 in lung tumors. After lung tumors were infected with SARS-CoV-2, viral titers in tumor homogenates were determined at 3, 15, and 30 dpi by plaque assay. Data are presented as mean ± s.d. **c,** Expression of SARS-CoV-2 N protein and S protein in tumor tissues. The mice were sacrificed at 30 dpi. Paraformaldehyde-fixed, paraffin-embedded tumor tissues were sliced to 5 µm thickness, and H&E staining and immunohistochemical staining were performed to detect SARS-CoV-2 S protein and N protein and hACE2. Scale bars, 25 µm. dpi, days post-infection; Anti-N mAb, anti-SARS-CoV-2 N protein monoclonal antibody; Anti-S Ab, anti-SARS-CoV-2 S protein polyclonal antibody.

**Supplementary Data Fig. 5.**
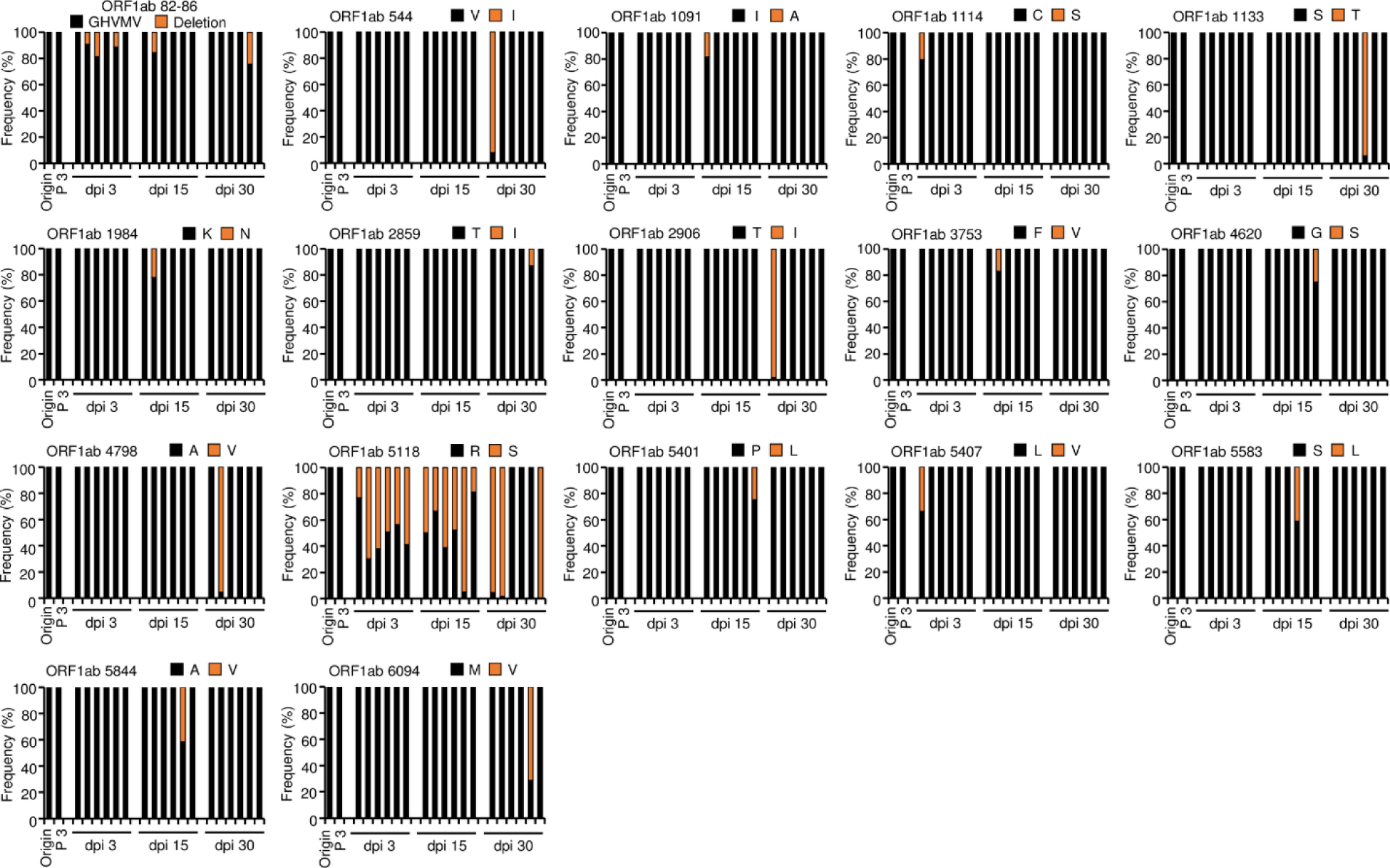
Frequencies and substitutions of amino acid sequences in wild-type SARS-CoV-2 ORF1ab during long-term infection in lung tumor tissues. Calu-3 cells were subcutaneously implanted into the right flank of NRGA mice (n = 6 per group). After the tumor volume reached 100 mm^3^, the tumors were inoculated with 1 × 10^6^ pfu of wild-type SARS-CoV-2. Whole-genome sequences of the viruses in the tumors were analyzed at 3, 15, and 30 dpi. Each panel indicates the accumulation and substitutions of mutations at an amino acid position in wild type SARS-CoV-2 during long-term infection in lung tumor tissues. Origin: amino acid sequences of wild-type SARS-CoV-2 (Supplementary Table 1). Passage 3 (P 3): amino acid sequences analyzed in wild-type SARS-CoV-2 after amplification in three passages in Vero E6 cells. dpi, days post-infection.

**Supplementary Data Fig. 6.**
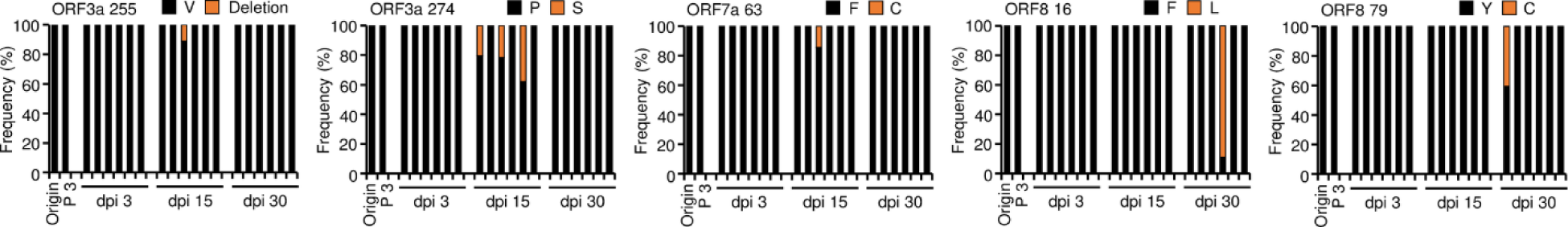
Frequencies and substitutions of amino acid sequences in wild-type SARS-CoV-2 ORF3a, ORF7a, and ORF8 during long-term infection in lung tumor tissues. Calu-3 cells were subcutaneously implanted into the right flank of NRGA mice (n = 6 per group). After the tumor volume reached 100 mm^3^, the tumors were inoculated with 1 × 10^6^ pfu of wild-type SARS-CoV-2. Whole-genome sequences of the viruses in the tumors were analyzed at 3, 15, and 30 dpi. Each panel indicates the accumulation and substitutions of mutations at an amino acid position in wild type SARS-CoV-2 during long-term infection in lung tumor tissues. Origin: amino acid sequences of wild-type SARS-CoV-2 (Supplementary Table 1). Passage 3 (P 3): amino acid sequences analyzed in wild-type SARS-CoV-2 after amplification in three passages in Vero E6 cells. dpi, days post-infection.

**Supplementary Data Table 1.**
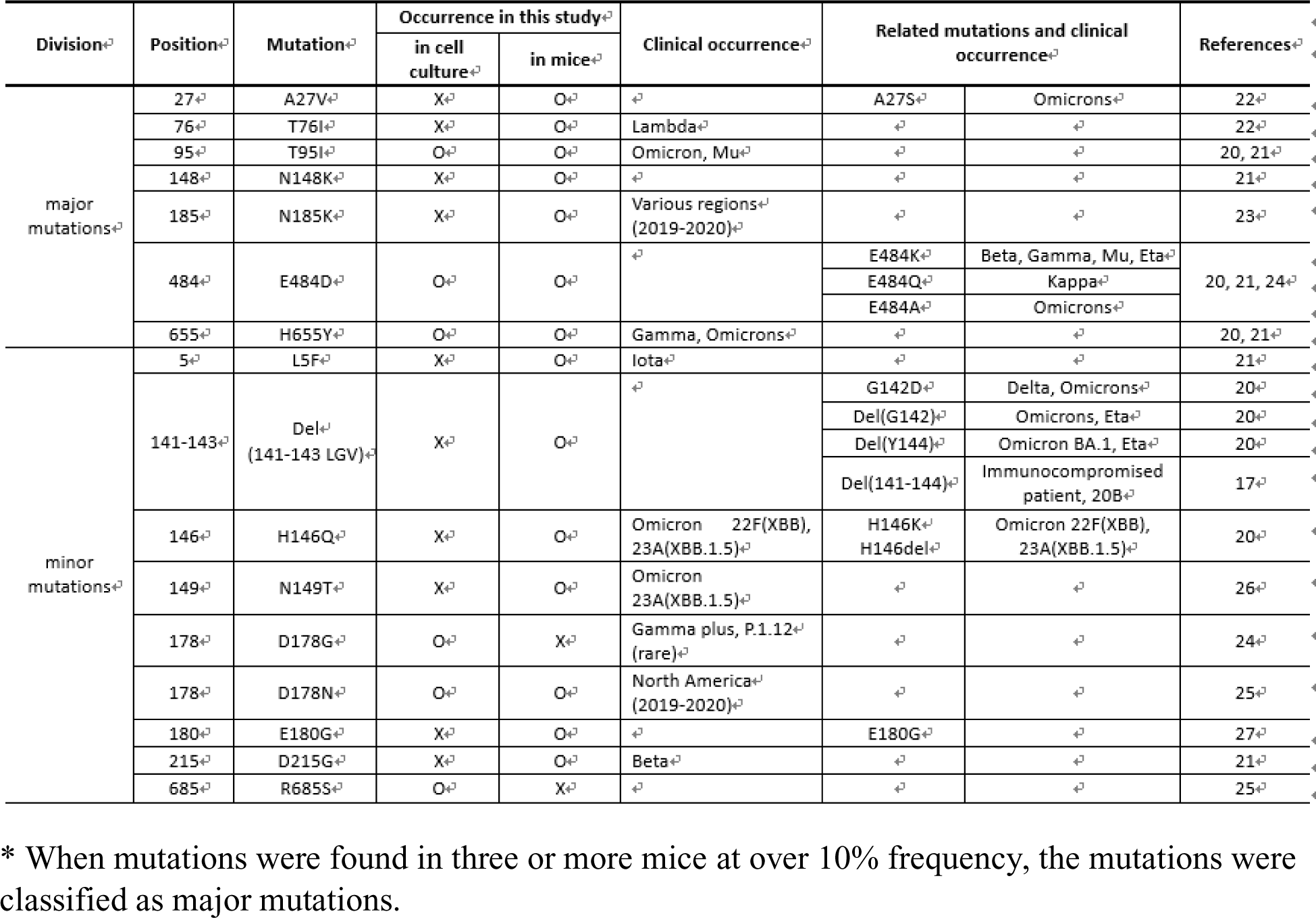
Summary of the S protein mutations found in this study * When mutations were found in three or more mice at over 10% frequency, the mutations were classified as major mutations.

**Supplementary Data Table 2.**
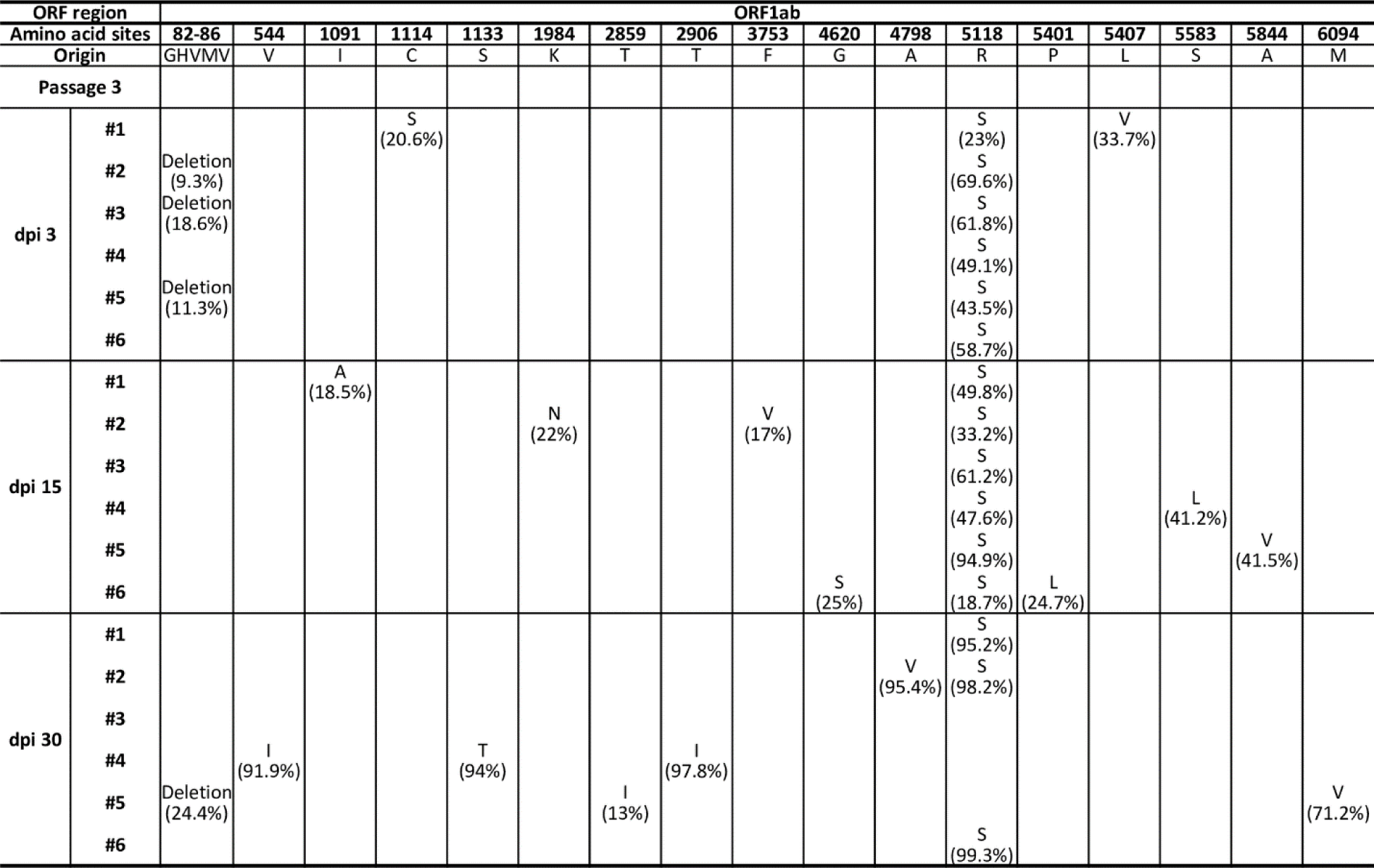
Frequency of mutations in wild-type SARS-CoV-2 during long-term infection in lung tumor tissues (ORF1ab).

**Supplementary Data Table 3.**
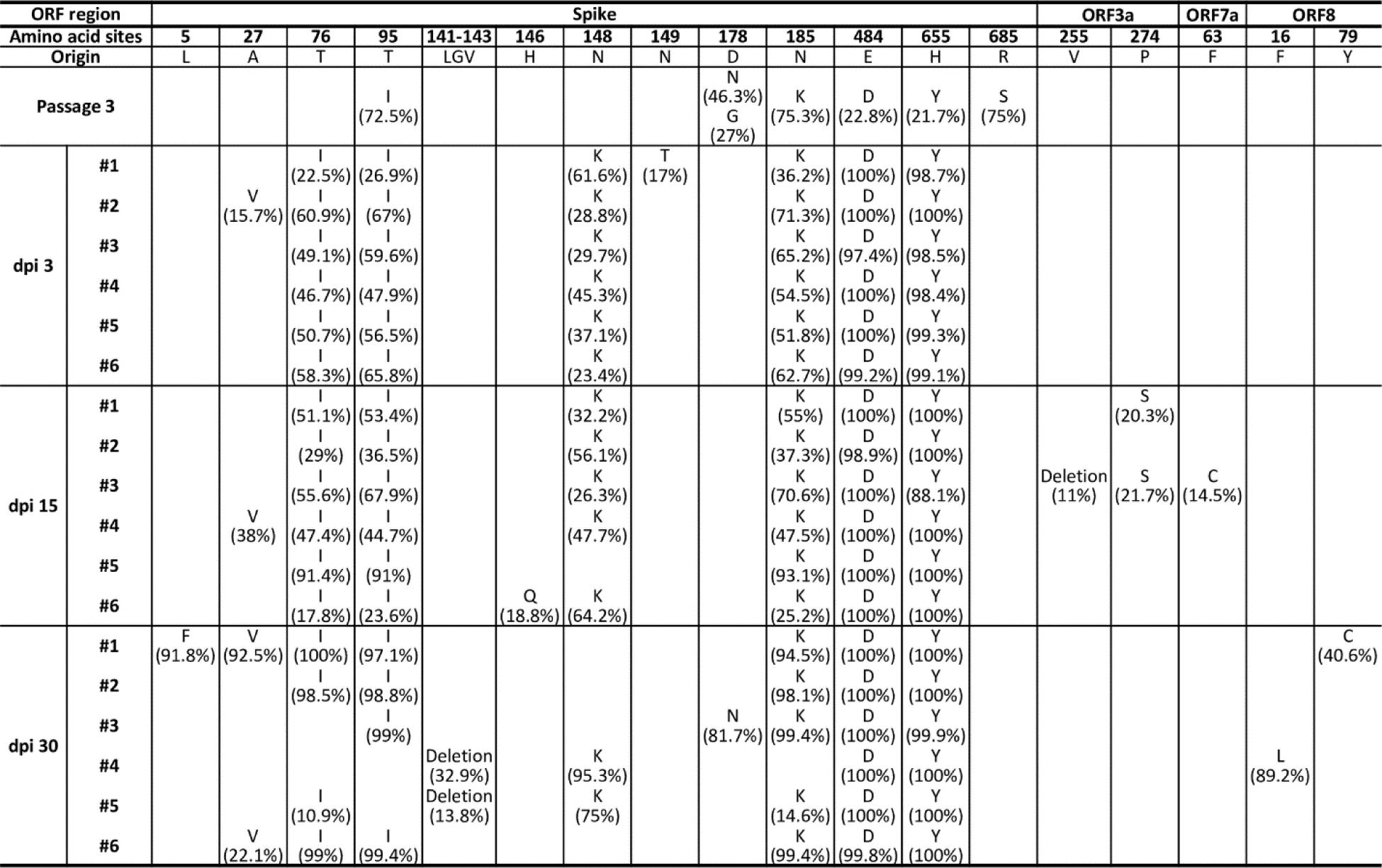
Frequency of mutations in wild-type SARS-CoV-2 during long-term infection in lung tumor tissues (Spike, ORF3a, ORF7a, and ORF8).

**Supplementary Table 1.** List of protein accession numbers of SARS-CoV-2 proteins based on GenBank: MW466791.1. (https://www.ncbi.nlm.nih.gov/protein?LinkName=nuccore_protein&from_uid=1955390757)

| Proteins | Amino acid no. | Accession no. |
| --- | --- | --- |
| ORF1ab | 7096 | QQO75521.1 |
| ORF1a | 4405 | QQO75522.1 |
| Spike | 1273 | QQO75523.1 |
| ORF3a | 275 | QQO75524.1 |
| Envelope | 75 | QQO75525.1 |
| Membrane | 222 | QQO75526.1 |
| ORF6 | 61 | QQO75527.1 |
| ORF7a | 121 | QQO75528.1 |
| ORF7b | 43 | QQO75529.1 |
| ORF8 | 121 | QQO75530.1 |
| Nucleocapsid | 419 | QQO75531.1 |
| ORF10 | 38 | QQO75532.1 |

